# Exploring genomic diversity and reproductive strategies in three expansion phases of the superdominant *Brachypodium rupestre* in high mountain grasslands

**DOI:** 10.1101/2025.06.05.657980

**Authors:** María Durán, Miguel Campos, Samira Ben-Menni, Alba Sotomayor, Pilar Catalán, Leticia San Emeterio, Rosa Maria Canals

## Abstract

Tall-grass expansion in natural plant communities is a concern under current global change scenarios. Driven by factors such as community vulnerability and the competitive abilities of tall-grass species, this phenomenon is exemplified by *Brachypodium rupestre* (Host) Roem. & Schult, a perennial tall-grass native to Europe. *B. rupestre* spreads aggressively in the grasslands of the western Pyrenees due to a disrupted regime of lack of herbivory and recurrent burnings that has persisted over several decades. This study aims to investigate whether the contrasted managements of grazing, abandonment and burnings have promoted specific reproductive traits and impacted the population genomics of *B. rupestre*. Nine sites varying in cover of *B. rupestre* classified into three expansion phases associated to different management regimes were monitored: i) scattered populations in multispecific grazed sites, ii) stand populations expanding at sites of relaxed grazing or abandonment, and iii) superdominant populations constituting dense degraded covers in recurrently burned sites. ddRADseq data showed high genomic diversity but relatively low genomic structure, likely due to substantial gene flow and the absence of wind barriers. The standardized index of association supported asexual reproduction in all populations, which is coupled with sexual reproduction. Contrasting management practices did not promote distinct genomic identities between expansion phases; rather, the genetic differentiation and divergence of populations responded to life history and spatial isolation, even within a small geographic area. The similar levels of genetic diversity and the significantly different number of clones between expansion phases suggest a complex pattern of genotypic and clonal variability, which may be influenced by environmental factors (historical disturbance regimes).

## 1 Introduction

Historical disturbance regimes of fire and herbivory have created and preserved open ecosystems worldwide, shaping its floristic composition through the promotion of grasses and impacting on their mechanisms of survival (Briske et al., 2005; Van Langevelde et al., 2003). These regimes have resulted in a balance between sexual and asexual regeneration strategies, promoting the survival of adults through regrowth or the promotion of offspring through seed production (Pausas & Keeley, 2014), which may have an impact on genetic variation within plant populations and their potential for expansion (Barrett et al., 2008; García-Ramos & Rodríguez, 2002; Schläpfer & Fischer, 1998).

Fire enhances the reproductive abilities of many annual and perennial grasses in contrasting Mediterranean (Canales et al., 1994; De Luis et al., 2005; Vidaller et al., 2018), temperate (Brys et al., 2005) and tropical environments (Angelo & Daehler, 2013; Damasceno & Fidelis, 2020) by improving spike and seed production, germination rates, and resprouting capacity. The location of buds and their capacity to survive fire events are crucial (Pausas & Paula, 2020), and asexual reproduction can benefit over sexual reproduction in many grasses. However, from a genetic perspective, recurrent fires may benefit sexual reproduction over regrowth due to the potential genetic load caused by the accumulation of deleterious somatic mutations over many fire cycles (Lamont & Wiens, 2003).

Regarding herbivory, it is the primary evolutionary force in the development of grasses, responsible for their ability to regrow through tillering (Gibson, 2009). Grasses combine sexual and resprouting mechanisms of persistence, with grazing playing a decisive role in determining the relative importance of these mechanisms through the timing and intensity of defoliation (Liston et al., 2003). Defoliation during vegetative growth enhances tillering and promotes regrowth over sexual reproduction when the apical meristem is eliminated (Gastal & Lemaire, 2015). The intensity of defoliation influences the balance between sexual and asexual reproduction. Optimal grazing pressure preserves basal meristems responsible for tillering and flowering, while high defoliation intensity may consume them.

Unbalanced fire and herbivory regimes can result in the degradation of chalk grasslands, threatening their high biodiversity (Durán et al., 2020; Tardella et al., 2020). In Europe, perennial tall-grasses of the *Brachypodium pinnatum* complex (*B. pinnatum* (L.) Beauv., *B. genuense* (DC.) Roem. & Schult.*, B. rupestre* (Host) Roem. & Schult) are common components of natural grasslands on chalk bedrock, constituting the cohort of other dominant grasses and composing very diverse montane communities (Canals & Sebastià, 2000). The decline of grazers and other factors can lead to aggressive expansion of these grasses (Bąba et al., 2012; Bobbink & Willems, 1987; Catorci et al., 2011; Múgica Azpilicueta et al., 2021). In the western Pyrenees, the expansion of *B. rupestre* is associated with reduced livestock farming and recurrent burning to manage late-season biomass accumulations (Canals et al., 2014). The species’ expansion depends on local management regimes, exhibiting varying phases. Under optimal grazing conditions, *B. rupestre* cover is low (<25%) developing in small, scattered clumps that constitute a component of grassland diversity. Decreasing grazing pressure leads to larger stands of ungrazed *B. rupestre* that progress within the grassland matrix. In ungrazed areas subject to recurrent burning, *B. rupestre* covers more than 70% of the soil surface and becomes a superdominant species, as defined by Pivello et al. (2018). This superdominance leads to the displacement of common grassland species and a consequent biodiversity loss.

The expansion of *B. rupestre* in a decoupled environment of low herbivory and high fire frequency is explained by a combination of factors such as large ecological versatility (Peralta, 2010), high tillering capacity and leaf plasticity (Bąba et al., 2016; Mojzes et al., 2003; Tardella et al., 2017), rapid response to nutrient pulses (Canals et al., 2014; Ryser et al., 1997), and early growth in spring and rapid loss of digestibility to herbivores (Catorci et al., 2014). Another important trait is the species’ vigorous rhizome, which transports nutrients to aboveground, stores toxic soluble aluminium (Bobbink et al., 1989; Canals et al., 2014; Pottier & Evette, 2010) and enables vegetative expansion through buried buds (Hamrick & Godt, 1996; Pausas et al., 2018; Pottier & Evette, 2010). *B. rupestre* is characterized by an outbreeding mating system (Catalan et al., 2016). Although diploid and tetraploid populations have been detected in the species (Schippmann, 1991), the more frequent cytotype is the allotetraploid (2n=4x=28), with individuals containing two different ancestral progenitor genomes of x=9 and x=5 chromosomes (Lusinska et al., 2019; Sancho et al., 2022). This allopolyploid origin may have influenced its hybrid vigour and ecological competitiveness (Soltis & Soltis, 2000). Population genomic studies based on genome-wide markers are appropriate methods for investigating the genetic diversity and structure of plant populations as well as for identifying patterns of variation, differentiation and genetic admixture. These studies provide information on population history and aid in our comprehension of their adaptive ecological processes (Luikart et al., 2019). They also serve to assess the genetic identity of individuals, reconstruct their genetic relationships, and group them into genetically separate clusters (Scariolo et al., 2021). Among the new methodologies used in population genomics, the ddRADseq approach produces high-density single nucleotide polymorphisms (SNPs) across the genome and provides data that can be efficiently analysed without the need for a reference genome sequence (Peterson et al., 2012). ddRADseq is also particularly suitable for the analysis of polyploid organisms because it enables the identification of loci that are shared across multiple subgenomes while minimizing issues related to assembly errors or homologous sequence misidentification (Ott et al., 2022). SNP-based multilocus genotypes can be also used to distinguish between sexually reproducing and clonally propagating asexual genotypes (Bailleul et al., 2016; Catalán et al., 2016). The extent of clonality can be large in rhizomatous grass populations and can affect their demographic and evolutionary trajectories. Theory predicts that the cost of outcrossing and reproductive assurance may lead to an over-representation of asexuals, which could eventually displace outcrossers from populations unless the latter are more likely to survive and reproduce (Holsinger, 2000).

The primary goal of this research is to examine the influence of contrasting disturbance mechanisms—grazing, abandonment, and fire—associated to different expansion phases on the genomic structure and diversity of neighbouring *B. rupestre* populations in grassland areas over the past century. Sexual reproductive traits of *B. rupestre* populations were analysed to assess investment in structures that promote outcrossing and genetic diversity. ddRADseq analyses were employed to determine genomic structure, clonality levels and genetic diversity of *B. rupestre*. Our initial hypothesis is that populations in sites exposed to different disturbance regimes and reflecting different expansion phases of *B. rupestre*, evolve towards different genomic constitutions. We consider that this divergence will be most evident in degraded *B. rupestre* grasslands that have experienced recurrent burning every 1-3 years for decades. We have tested this hypothesis by monitoring *B. rupestre* individuals at different phases of community degradation, from diverse grasslands cohorts to primary constituents of monophyte covers.

## 2 Materials and methods

### 2.1 Study area and sampling sites

The study was developed in western Pyrenees and covered an area of natural grasslands located at the head of the neighbouring valleys of Aezkoa (Spain) and Cize (France) at altitudes ranging 800 and 1400 m.a.s.l. The area is protected by the Natura 2000 network and includes the SAC *Roncesvalles-Selva de Irati* (ES0000126) and the SAC *Pic d’Herrozate et foret d’Orion* (FR7212015). The climate is snowy and cold in winter and temperate and foggy in summer. According to the Irabia climatic station, the average annual temperature and precipitation are 9.4 °C and 1881.2 mm, respectively. The predominant wind direction is from the north-west (http://meteo.navarra.es). The floristic communities include beech forests (*Fagus sylvatica*), heathlands (*Erica vagans*, *Calluna vulgaris*), gorselands (*Ulex gallii*) and natural grasslands constituted by perennial grasses (such as *Festuca* gr. *rubra*, *Agrostis capillaris*, *Danthonia decumbens*, *Brachypodium rupestre*), forbs (such as *Achillea millefolium*, *Potentilla erecta*, *Gallium saxatile*) and legumes (such as *Trifolium repens* and *Lotus corniculatus*).

Despite current geopolitical boundaries, a historic treaty regulates the joint use of grasslands by Spanish and French livestock and an extensive mixed ranging of sheep, cattle and horses occurs from May to October (Razquin et al., 2012). Pastoral burns are a traditional tool used to complement herbivory and reduce shrub encroachment and are applied in small patches (shrub-to-shrub) with low recurrence (> 7 years). However, the sharp decrease in livestock loads in the last century has caused in certain areas large accumulations of vegetation that are reduced by increasing the intensity and the recurrence (1-3 years) of burns (Canals, 2019; Múgica et al., 2021). Based on previous research, we selected 9 sampling sites that differed in *B. rupestre* cover and disturbance regime, encompassing a variety of situations of herbivore pressure and fire recurrence (Ferrer & Canals, 2008). *B. rupestre* populations from the sampling sites were classified according to their expansion phase as follows: 1) scattered populations (S), when *B. rupestre* was a constituent of multispecies grasslands (average cover < 25%) in optimally grazed and not frequently burned areas, 2) stand populations (P) when *B. rupestre* grew mostly in stands that covered between 25% and 70% of the surface in low or abandoned grazing areas, and 3) superdominant populations (I) when *B. rupestre* exceed 70% of cover and was evenly distributed in areas ungrazed and frequently burned (Table 1, Figures 1 and 2).

**Figure 1.**
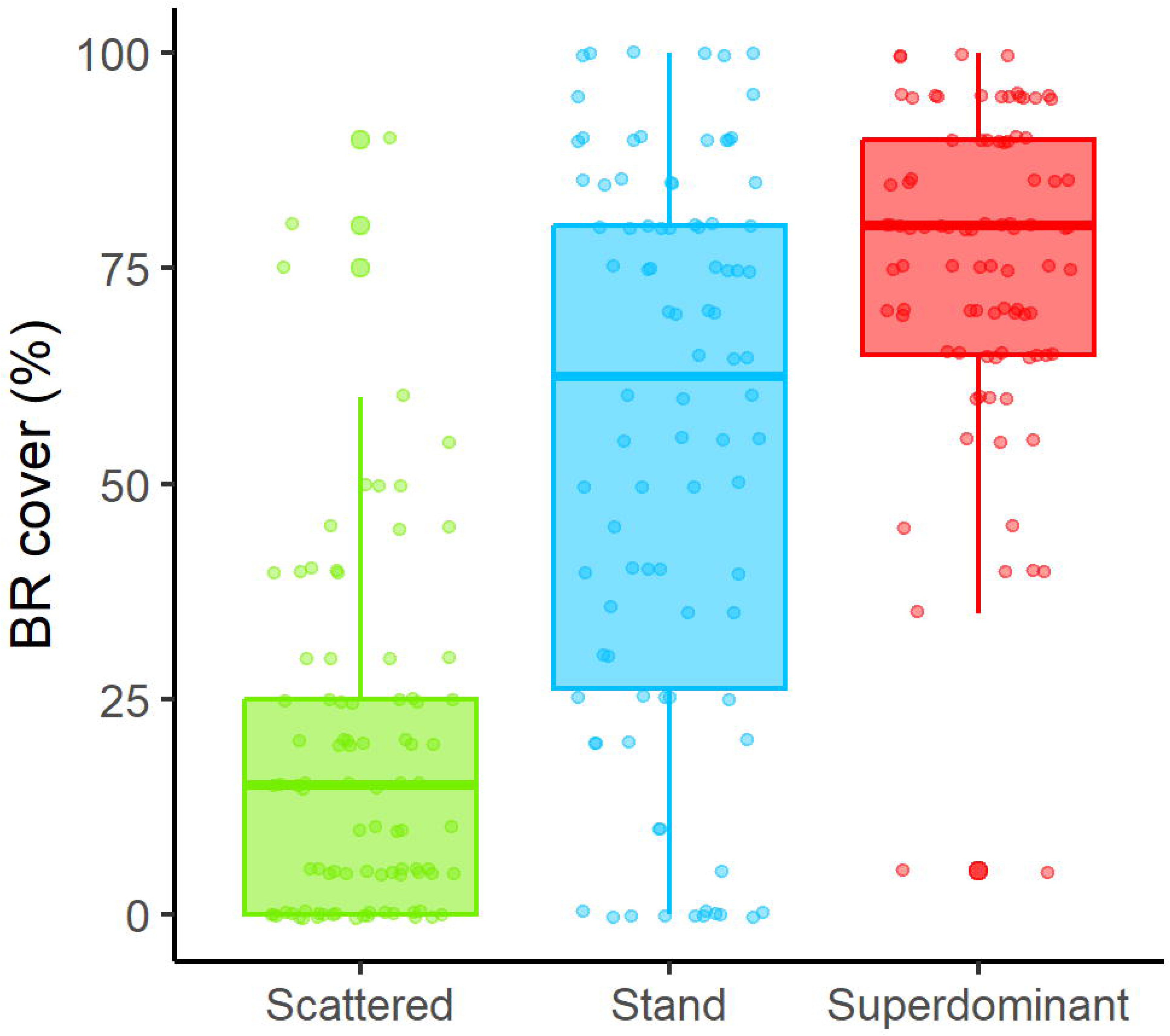
Percentages of cover of *B. rupestre* in the different expansion phases in the study area.

**Figure 2.**
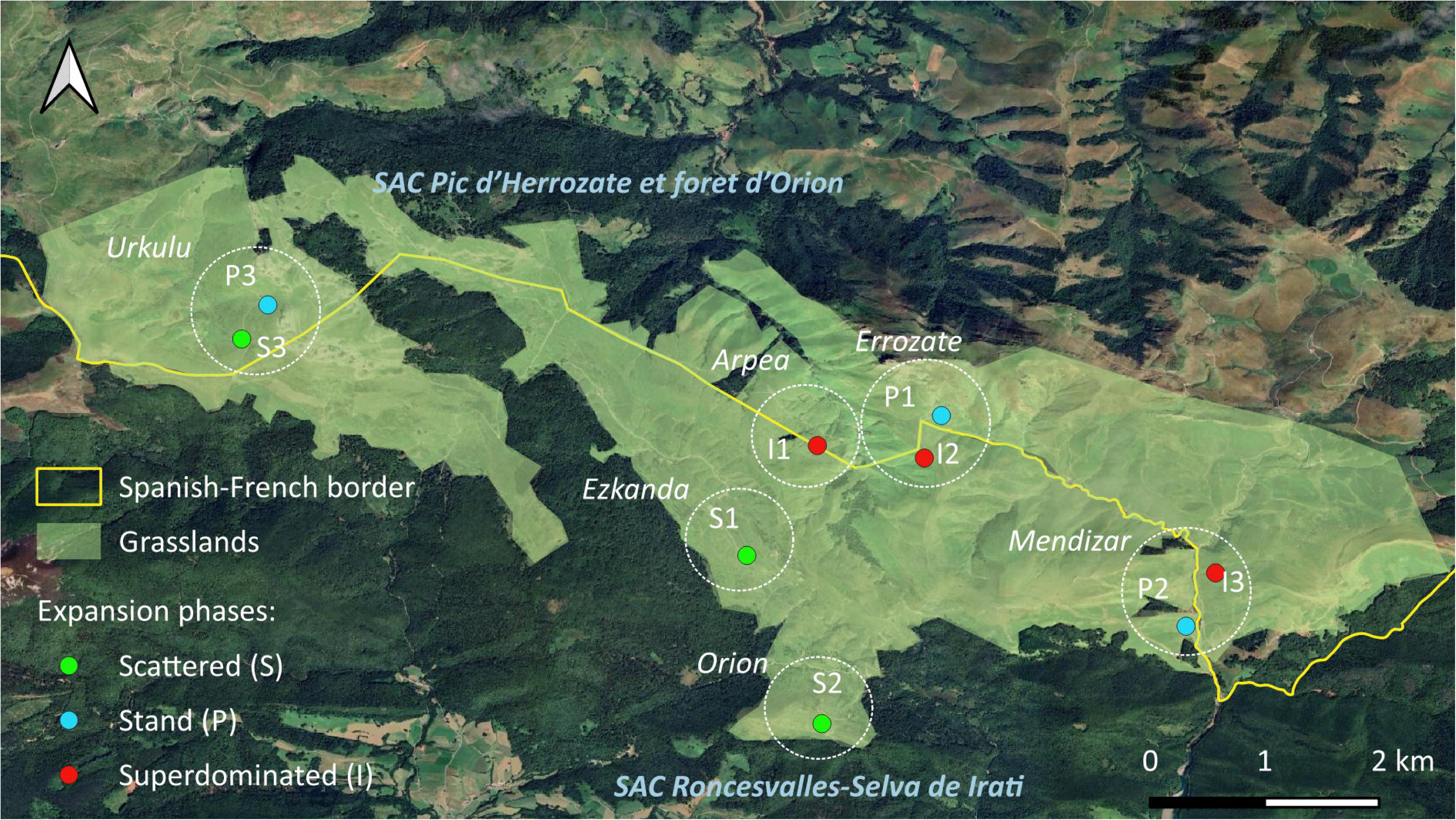
Location of the study area in the western Pyrenees and detail of the six zones (dotted circles) that include the nine sampled sites in different expansion phases: Scattered (S): <25%; Stand (P): 25-70%; Superdominant (I): >70%.

**Table 1.** Sampling sites classified according to the *B. rupestre* expansion phases (scattered, stand and superdominant), fire and grazing management and topographic characteristics.

|  | Sites | Expansion phase | Management |  | Topographic characteristics |  |  |
| --- | --- | --- | --- | --- | --- | --- | --- |
|  |  |  | Grazing | Burnings | Altitude (masl) | Slope (%) | Aspect |
| 1 | Ezkanda-S1 | Scattered | Grazing | > 7 years | 1060 | 10 | SW |
| 2 | Orion-S2 |  | Grazing | > 7 years | 1070 | 10 | NW |
| 3 | Urkulu-S3 |  | Grazing | > 7 years | 1290 | 30 | SW |
| 4 | Errozate-P1 | Stand | Abandonment | > 7 years | 1090 | 50 | E |
| 5 | Mendiziar-P2 |  | Grazing | > 7 years | 860 | 30 | E |
| 6 | Urkulu-P3 |  | Abandonment | > 7 years | 1290 | 30 | SW |
| 7 | Arpea-I1 | Super dominant | Abandonment | 1-2 years | 940 | 50 | NE |
| 8 | Errozate-I2 |  | Abandonment | 1-2 years | 1090 | 50 | W |
| 9 | Mendiziar-I3 |  | Abandonment | 1-2 years | 860 | 30 | E |

### 2.2 Experimental field sampling

In June 2020, we established a 141 m transect corresponding to the hypotenuse of a rectangle with sides of 100 x 100 m (1 ha area) in each sampled population (Table S1). Along the hypotenuse line, thirty quadrats of 1 × 1 m were distributed in 3 parallel rows following the design of Figure 3, box A. In each quadrat, the cover of *B. rupestre* was estimated and the panicles of all individuals were counted, collected and transported to the Public University of Navarre laboratory where all seeds were extracted and counted. Fifty seeds from each quadrat were placed in petri dishes with moist Whatman paper, except in those quadrats with less than fifty seeds, in which all the collected seeds were used. Petri dishes were checked daily and germination rates were calculated.

**Figure 3:**
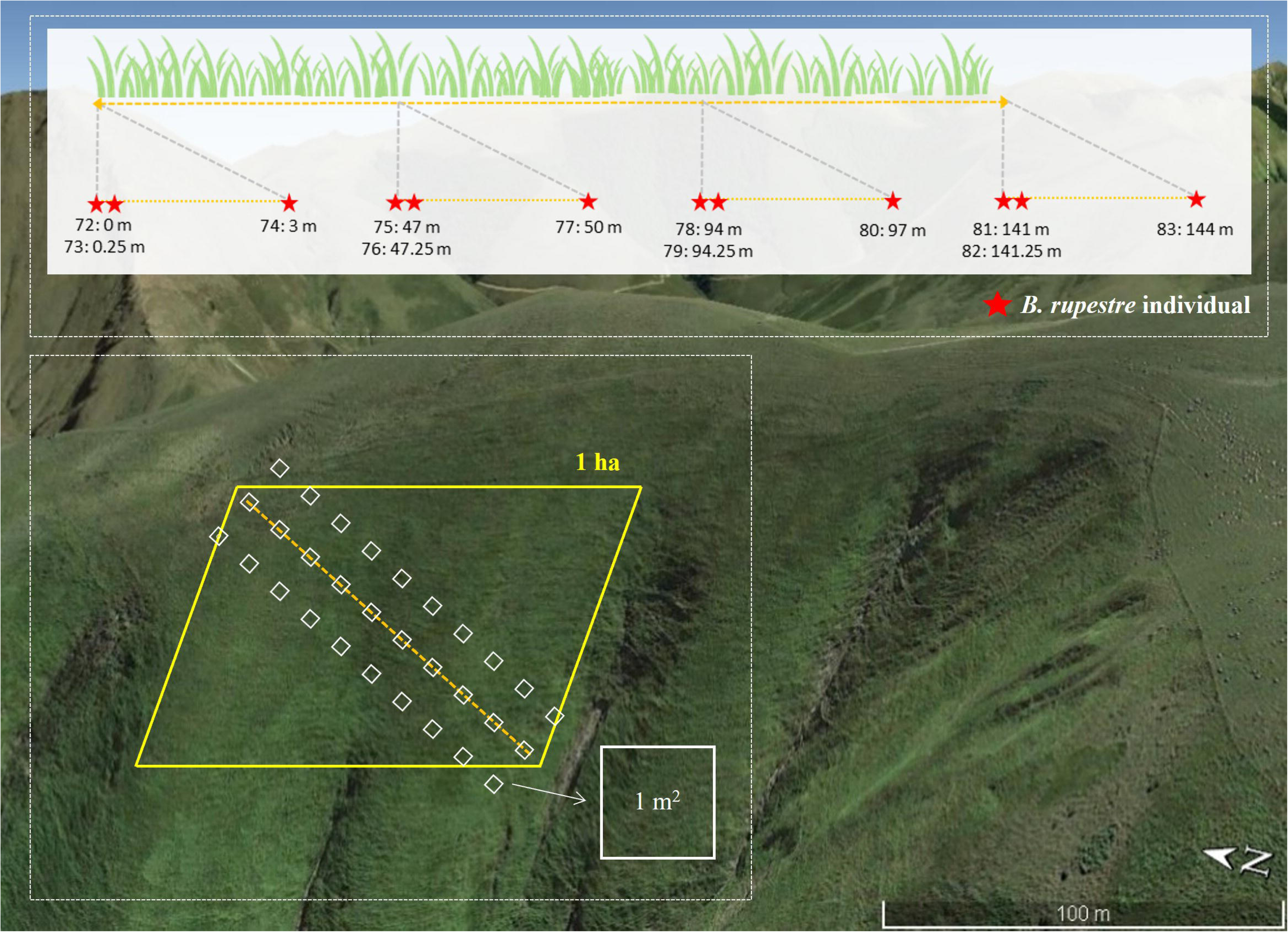
Example of sampling design in Errozate-I2. Collections of *B. rupestre* panicles for reproductive analysis (box A) and of *B. rupestre* plants for genetic analysis (box B).

Additionally, nine to twelve plants were collected in each sampled site at different distances from each other (Figure 3, box B; Table 2). Individuals were collected along a transect line in groups of three at the beginning (0, 0.25 and 3 m), in the middle (47, 47.25 and 50 m) and at the end (94, 94.25 and 97 m). In certain transects, we collected an additional set of three individuals at 141, 141.25 and 144 m. As a result, for each transect we had between 3 and 4 pairs of individuals collected 0.25 m apart and between 9 and 12 pairs of individuals collected less than 3 m apart. Plants collected close to each other (0.25 cm distance) were more likely to be clones, while those located at greater distances (3 m – 47 m) increased the probability of sampling different genets (Bąba et al., 2012). In total, fresh and healthy leaves of *B. rupestre* belonging to 95 individuals were collected and processed for DNA extraction. The taxonomic identity of all the individuals used in the study was confirmed through morphological analysis following Schippmann (1991).

**Table 2.** Sample size for reproductive and genomic analyses of *B. rupestre*. Samplings were done in plots of 30 m^2^.

|  | Sites | Expansion phase | Number of individuals for genomic analysis | Number of individuals (range) for reproductive analysis |
| --- | --- | --- | --- | --- |
| 1 | Ezkanda-S1 | Scattered | 7 | [1 - 9] |
| 2 | Orion-S2 |  | 10 | [10 - 19] |
| 3 | Urkulu-S3 |  | 10 | [20 - 29] |
| 4 | Errozate-P1 | Stand | 10 | [30 - 39] |
| 5 | Mendiziar-P2 |  | 9 | [40 - 49] |
| 6 | Urkulu-P3 |  | 9 | [50 - 59] |
| 7 | Arpea-I1 | Super dominant | 11 | [60 - 71] |
| 8 | Errozate-I2 |  | 12 | [72 - 83] |
| 9 | Mendiziar-I3 |  | 12 | [84 - 95] |

### 2.3 Data analysis of reproductive data

Seed production (n seeds m^-2^) was calculated counting the total number of seeds produced in the 1 m^-2^ subplots. The relationships between seed production and *B. rupestre* cover and the expansion phase were analysed using generalized additive models for location, scale and shape (GAMLSS) with the gamlss package (Rigby & Stasinopoulos, 2005). GAMLSS is a distribution-based semiparametric regression approach capable of fitting up to four parameters of the data distribution [location (e.g. mean), scale (e.g. variance) and shape (skewness and kurtosis)]. GAMLSS allows choosing from a wide variety of family distributions for seed counts and all parameters of the distributions can be modelled as functions of the explanatory variables. The distribution of seed counts was positively skewed and zero-inflated, and the relationship between seed counts and *B. rupestre* cover was non-linear. For these reasons, we fitted a model with a zero-inflated negative binomial distribution (ZINBI) using seed production as the response variable, and the expansion phase (scattered, stand and superdominant populations) and a smoother cubic spline of *B. rupestre* cover as explanatory variables. The phase of expansion of *B. rupestre* is associated with management variables (burning and grazing see Table 1). The scattered populations are associated with grazing and low burning frequency, the stand populations are associated with grazing abandonment and low burning frequency and the superdominant populations are associated with grazing abandonment and high burning frequency. We used likelihood ratio tests to choose the best model with the following strategy: 1) We selected the explanatory variables for the mean (μ) using a forward approach, 2) we selected the explanatory variables for the dispersion parameter (σ) using a forward approach, 3) we selected the explanatory variables for the probability of inflation at zero (ν) using a forward approach, 4) we checked if the terms for σ were necessary using a backwards approach, 5) we checked if the terms for μ were necessary following a backwards approach, and 6) we checked whether a random intercept was necessary for each site. Finally, we tested whether other distribution families for seed counts produced a best fit. We compared the number of seeds per inflorescence and the germination percentage among expansion phases estimating 95% confidence intervals using bootstrap with replacement (10,000 resamples).

### 2.4 Ploidy level estimations using genome sizes and site-based heterozygozities

Samples from 90 individuals of *B. rupestre* were used in the genomic study. The genome size (2C value) of six representative samples of individuals from the Arpea (3) and Urkulu (3) populations (Figure 2) was estimated using propidium iodide staining of cell nuclei and flow cytometry measurements (Sysmex Ploidy Analyser) according to Doležel et al., (2007). The final DNA content of each individual was calculated based on two independent measurements (on at least 5000 nuclei each), which showed a coefficient of variation < 3, and using *Solanum lycopersicum* L. “Stupické polní rané” (1.96 pg/2C) as internal standard. Additional ploidy level estimates were performed for the total number of sequenced individuals under study (90) using site-based heterozygosity with the nQuack program (Gaynor et al., 2024).

### 2.5 ddRADseq genomic libraries and population genomic analysis using SNPs genotyping

Genomic DNA was extracted from fresh tissues using an adjusted CTAB method (Doyle & Doyle, 1987; Murray & Thompson, 1980). The DNA quality and concentration of each individual sample were measured with a Qubit analyser (Life Technologies Corporation, Caribad, CA). After DNA extraction, samples were submitted to Floragenex (https://www.floragenex.com/) for construction of ddRADseq libraries and sequencing, following Truong et al. (2012). The ddRAD sequence data were analysed using Ipyrad software (Eaton & Overcast, 2020) (Table S2). Quality filtered pair-end raw reads were demultiplexed via barcoding, trimmed for adapter contamination, and loci were assembled by *de novo* clustering due to the lack of a reference genome. Assembly was performed using the following Ipyrad parameters: max_low-qual 0, phred_Qscore 33, mindepth 6, maxdepth 10000, clust-thresold 0.85, max-barcode_mismatch 0, max_alleles_consens 2, max_Ns_consens 0.05, min-samples_locus 5, max-SNPs_locus 0.2, max_Indels_locus 8, max-shared_Hs_locus 0.5 (https://github.com/Bioflora/Brupestre_ddRAD). A total of 153,176 loci were present in the studied samples. To decrease potential linkage-disequilibrium (LD) bias, we applied LD filtering using PLINK2 with window size of 500 kb, a step size of 5 variants, minimum allele frequency (MAF) set in 0.05, and a threshold of 0.2 to identify and exclude linked SNPs and singletons. Accession ddRADseq data used in this study were deposited in the European Nucleotide Archive (ENA) (Table S2).

An unrooted maximum likelihood (ML) phylogenomic tree was constructed for all the studied individuals using the concatenated SNP data set through the IQtree program (Nguyen et al., 2015), imposing a best-fitting GTR+GAMMA substitution model and 1,000 bootstrap replicates. A Bayesian model-based analysis was performed to infer the genomic structure of *B. rupestre* populations and the spatial genomic ranges of *B. rupestre* using ADMIXTURE v.1.3.0 (Alexander et al., 2009). This software alternately updates the allele frequency and ancestry fraction parameters by using a block relaxation approach that retrieves data from the multilocus SNP genotype data matrix. One to ten hypothetical genetic groups (*K*=1-10), corresponding to the number of sampled populations plus one, were tested with ten iterations for each *K*. We used a cross-validation procedure to choose the best *K* for our data. For each run, we obtained a cross-validation (CV) error reported in the result, and the best *K* was selected based in its lowest CV error values (Alexander et al., 2009). Admixture results for each K value were plotted using the Pophelper package in R (Francis, 2017). The best *K* histogram was added to the ML phylogenomic tree using the online tool iTOL (Letunic & Bork, 2021). In order to estimate the robustness of the population genomic structure, we additionally calculated pairwise Fst genetic distances between populations using the dartR package in R (Mijangos et al., 2022).. To test for significant partitioning of the variance of alternative population genetic structures that would better explain the observed genomic data, we conducted an analysis of molecular variance (AMOVA) with Arlequin v.3.5.2.2 (Excoffier et al., 1992). We performed a standard AMOVA with all populations and also tested four different scenarios through hierarchical AMOVAS, grouping individuals by geographic zones, expansion phase (S, P, I), and grazing and burning management treatments. We ran the AMOVAs with 10,000 permutations to quantify the variance between groups.

### 2.6 Genomic diversity of populations and clonal detection

Genomic diversity was assessed within populations and among population groups. We estimated the observed heterozygosity (Ho), the expected heterozygosity (Hs), and the inbreeding coefficient (FIS) of populations with Plink2 (Chang et al., 2015), while the rate of selfing (s) was calculated as s=2Fis/(1+Fis) (Ritland, 1990). Significantly higher values of observed heterozygosity than expected suggested gene flow among populations, while lower values suggested genetic isolation and inbreeding.

We followed Bailleul et al. (2016) and Schuler et al. (2021) for the detection of putative intrapopulation genetic clones and the analysis of clonality descriptors. The concept of multilocus lineages (MLL) is used to define clusters of multilocus genotypes (MLGs) that belong to the same genet, and therefore share the same original event of sexual reproduction but are slightly different due to somatic mutations (Bailleul et al., 2016). To define the potential existence of clonal lineages or MLLs within populations (i.e., different multiple locus genotypes (MLGs) belonging to the same clone), we analysed the distribution of frequencies of genetic distances between pairs of MLGs using Bruvo distances implemented in the POPPR program (Kamvar et al., 2014). We calculated the following clonality descriptors to characterise the clonal richness of the populations: number of MLLs, number of expected MLLs (eMLLs), and clonal richness (R) corrected for sampling size, defined as the number of multilocus genotypes (MLG) relative to the number (N) of samples assessed R=(MLG–1)/(N–1) (Dorken & Eckert, 2001), and their genotype diversity: Shannon-Weiner Diversity index (H), Simpson’s index (lambda; corrected for sampling size), and clonal evenness index (E.5), which shows how equally each MLL is represented. We also calculated the standardized association index (rd) (Agapow & Burt, 2001) to test for the predominant reproductive model (sexual *vs.* clonal) with values differing significantly from zero being indicative of clonal propagation. The significance of rd was tested through 1,000 permutations.

## 3 Results

### 3.1 Numbers of inflorescences and seeds and germination rates

A total of 4,139 seeds of *B. rupestre* were extracted from 2,360 inflorescences collected in a total area of 270 m^2^ (Table 3). The model selected for seed counts included the cubic spline of *B. rupestre* cover with three degrees of freedom, the expansion phase and their interaction term as explanatory variables for the local parameter (μ), and the cover of *B. rupestre* for the shape parameter (ν) (Table 4). Seed counts per unit area (m^2^) varied significantly among expansion phases (LRT=6.6, p=0.037, Table 4), being lowest in in the scattered populations and reaching a maximum in the stand populations (Figure 4 top left). Similar patterns were found when considering seed count per inflorescence (Figure 4, bottom left). Germination percentages did not differ among expansion phases (Figure 4, bottom right).

**Figure 4.**
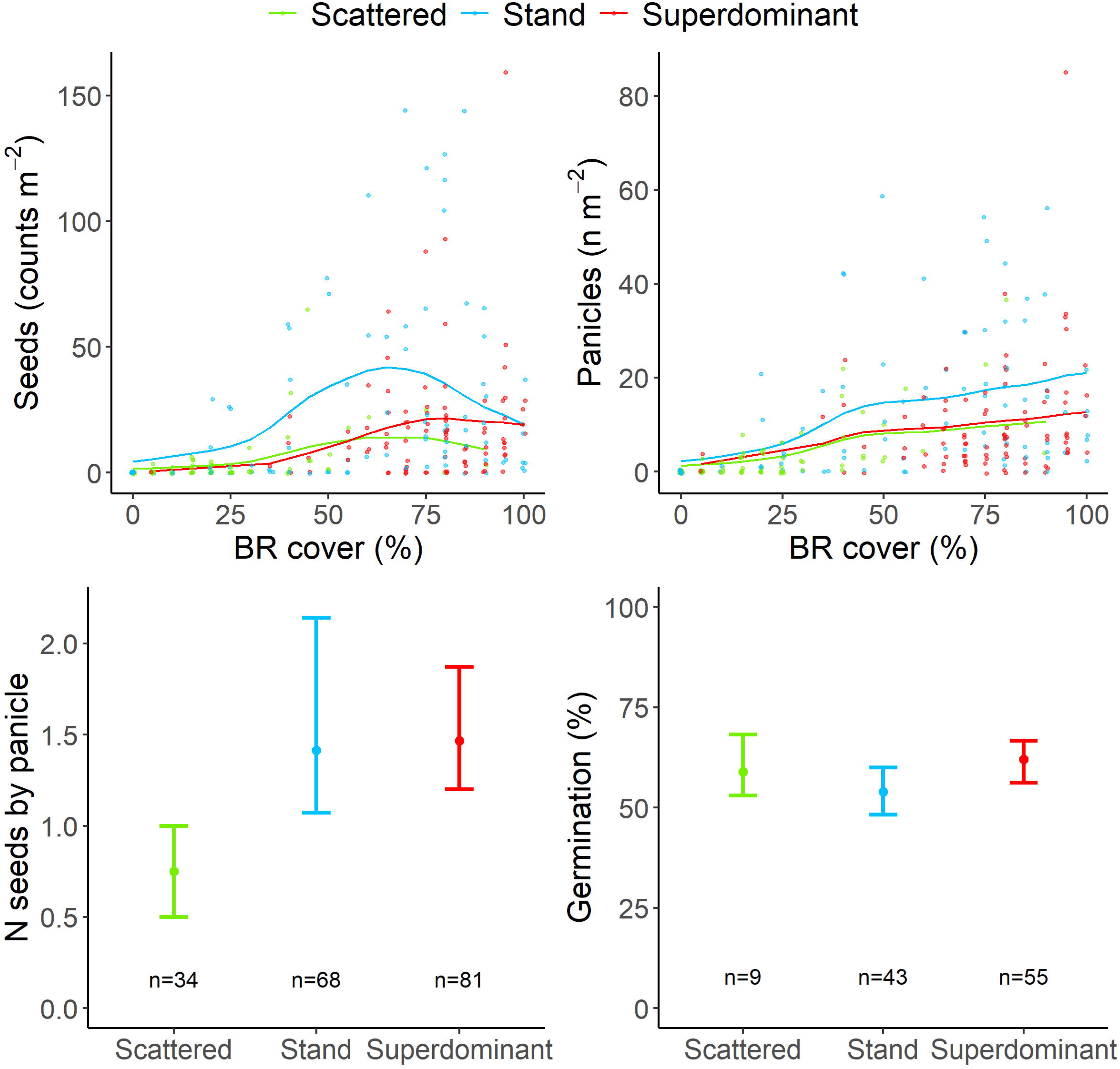
GAMLSS model fitted values for seeds counts (top left) and inflorescences counts (top right) as function of *B. rupestre* cover and expansion phase (S, P, I). Points, raw data; lines, fitted values. Median and 95% confidence intervals of number of seeds per inflorescence (bottom left) and seed germination percentage (bottom right) for the *B. rupestre* populations at different expansion phases (S, P, I).

**Table 3:**
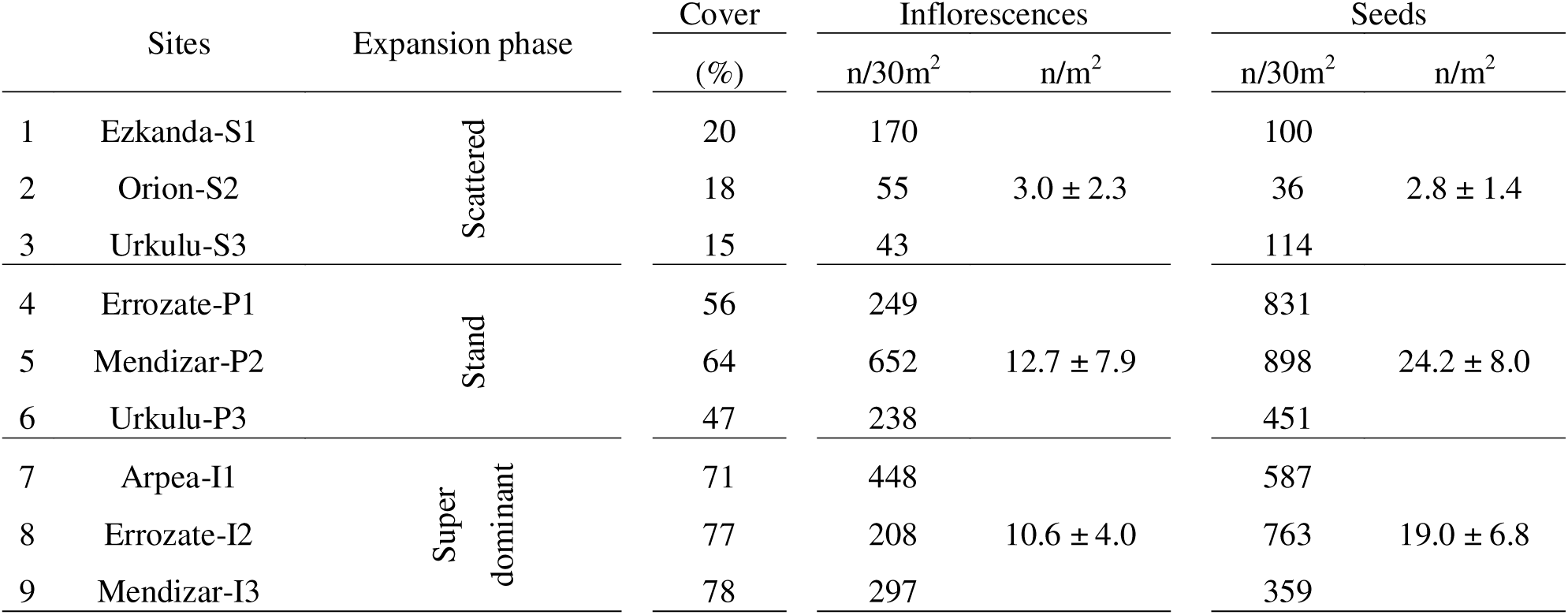
Inflorescences and seed counts of the studied *B. rupestre* populations.

**Table 4.** Likelihood ratio tests (LRT) for the significance of individual terms of the GAMLSS model for seed and inflorescence counts. Local (μ) and shape (ν) parameters.

| Parameter | Explanatory variable | LRT | p |
| --- | --- | --- | --- |
| Inflorescence |  |  |  |
| $\mu$ | cs(Cover) | 161.3 | <0.001 |
|  | Expansion phase | 10.7 | 0.005 |
|  | cs(Cover) : Expansion phase | 2.5 | 0.289 |
| $v$ | Cs(Cover) | 21.8 | <0.001 |
| Seeds |  |  |  |
| $\mu$ | cs(Cover) | 141.3 | <0.001 |
|  | Expansion phase | 12.4 | 0.002 |
|  | cs(Cover) : Expansion phase | 3.8 | 0.145 |
| $v$ | Cover | 19.9 | <0.001 |

### 3.2 Ploidy level assignation

The flow cytometry genome size analysis results indicated an average content of, respectively, 1.401 to 1.410 2C/pg for Arpea and Urkulu individuals (Table S3) that corresponded to allotetraploid plants of *B. rupestre* (Sancho et al., 2022). For the site-based heterozygosity analyses of the 90 individuals under study with nQuack we first selected the “normal-uniform fixed model” as the best model to accurately estimate the ploidy level of *Brachypodium* samples (Table S4a), using as controls RADseq data from samples of known ploidy, such as those of the diploids *Brachypodium stacei* (2) and *B. distachyon* (2) (Campos et al., 2024), and those of allotetraploid *B. rupestre* [the Arpea (2) and Urkulu (2) individuals mentioned above] (Table S3). We then applied this model to all the *B. rupestre* samples studied, imposing a search of 1000 bootstrap replicates, and obtained a tetraploid assignation for all of them (Table S4b). Allotetraploid *B. rupestre* is a diploidized amphidiploid (2n=4x=28; x=9+5) and therefore behaves cytologically and genetically as a functional diploid (Catalán et al., 2016; Sancho et al., 2022; Schippmann, 1991). Allopolyploids that behave like perfect amphidiploids could be analyzed as double diploids, using the genetic parameters described for diploid organisms (Catalán et al., 2006; Weir & Cockerham, 1984).

### 3.3 ddRADseq data and population genomics

Sequencing of the ddRADseq libraries generated over 447 million PE-reads (Table S2) with an average of 4.71 million PE-reads per sample. After data quality filtering, an average of 77,446 96bp-length reads per *B. rupestre* individual were used for the final assembly (Min: 32,772, Max: 90,461). Between 15,561 and 62,588 loci were retained in the individual samples after filtering for shared presence threshold among genotyped individuals. Specifically, loci were retained if they were present in at least 5 samples, as specified by the min_samples_locus parameter in the Ipyrad pipeline. Five individuals with low quality sequences (Mendizar-P2_45, Urkulu-P3_56, Arpea-I1_70 and Ezkanda-S1_1 and 7) were removed from the final analyses. An initial data matrix of 378,959 SNPs, including conserved invariant positions, was constructed for the remaining 90 individuals. A final comprehensive data set of 72,973 SNPs resulted from filtering of SNPs showing linkage disequilibrium (LD) among genotyped individuals with MAF > 0.05, excluding singletons and invariant positions, and with 67.76% missing data.

The ML phylogenomic tree revealed strongly supported divergences for the main populations and subpopulations linages of *B. rupestre* in Aezkoa, which showed bootstrap support (BS) values >83% for the main lineages and a strong topological concordance with the geographic distribution (Figures 5, 6). The more spatially distant western Urkulu populations nested within a highly supported clade (94% bootstrap support (BS) that separated from the rest and further split into two sister subclades, the Urkulu-P3 and the (predominantly) Urkulu-S3 subpopulations (Figure 6). The second group (100% BS) branched into two sister clades. The first clade (83% BS) comprised two weakly supported subgroups including samples from the centrally distributed populations of Arpea I1 (plus three individuals from the nearby populations of Ezkanda (2) and Orion (1)) (45% BS) and Ezkanda (plus two individuals from the nearby population of Arpea) (44% BS). The second strongly supported clade (100% BS) recovered the twinning relationship of the southern population of Orion S2 (plus six individuals from Errozate P1) (85% BS) with a clade (84% BS) consisting of central-eastern populations of Errozate (72% BS) and Mendizar (plus three individuals from Orion) (66% BS). The Errozate clade was divided into three subpopulation subclades (Errozate-I2/(Errozate-P1, Errozate-I2)) and the Mendizar clade into two subpopulation subclades Mendizar-P2/Mendizar-I3 (plus Orion-S2), which had relatively high support (Figure 6).

**Figure 5.**
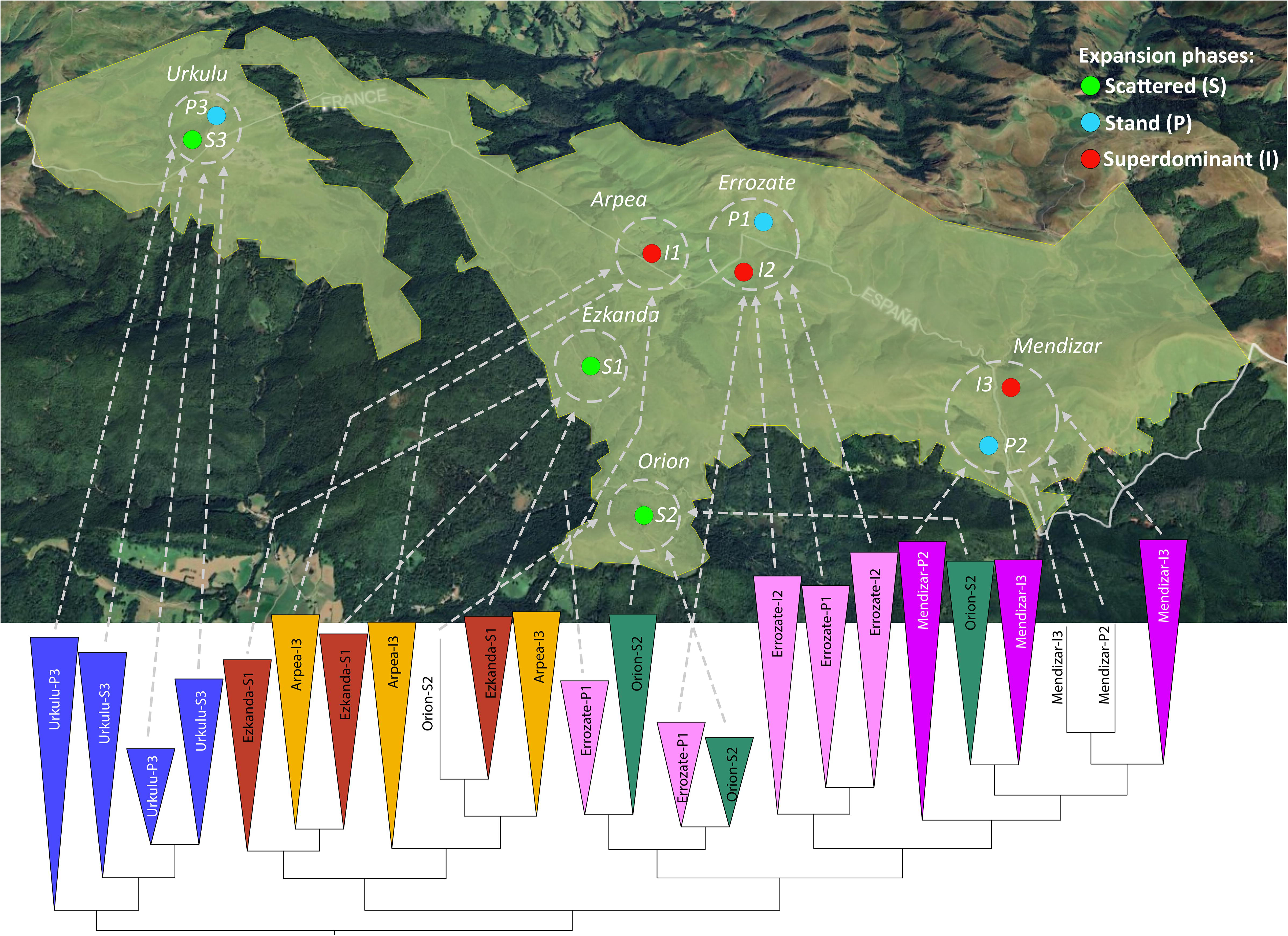
Summarized ML phylogenomic tree showing the concordance of the topological resolution of the *B. rupestre* population lineages and their respective West-to-East geographic distribution. Colour codes of expansion phases are indicated in the chart.

**Figure 6.**
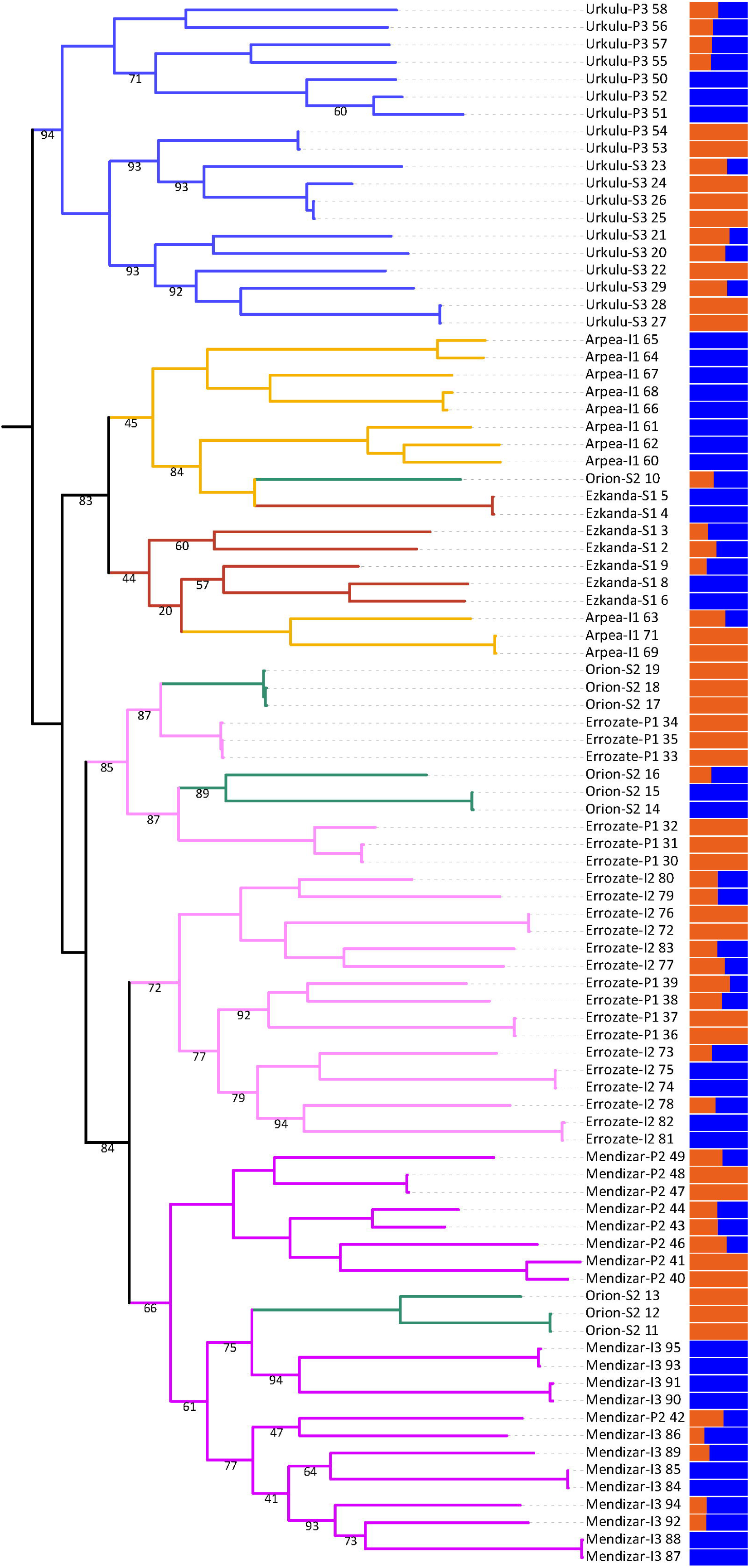
Phylogenomic tree and genomic structure of the sampled *B. rupestre* populations. (A) Maximum likelihood tree constructed using 90 individual samples and 72,973 SNPs with IQtree. Percentages of bootstrap support values (BS) <95% are indicated below branches; the remaining branches have BS=100%. Colour codes of branches correspond to populations indicated in Figure 5. (B) Graph with coloured bars showing the percentages of genomic membership of the samples to the K=2 optimal hypothetical genomic groups obtained with ADMIXTURE (cluster 1, blue; cluster 2, orange).

The genomic structure detected by ADMIXTURE indicated that the most suitable clustering for the data was K=2, based on the lowest cross-validation error (CV_error_=0.884), while the other K values showed significantly higher values (Figure S1). The structure revealed for the best two groups did not fully align with the expected populations and geographical clusters observed in the ML phylogenomic tree (Figure 6), nor with those of the other K suboptimal groups (Figure S1). However, K=2 results indicate that individuals from Urkulu-P3, Arpea-I1, Ezkanda-S1, and Mendizar-I3 primarily form cluster 1, while those from Urkulu-S3, Orion-S2 and Mendizar-P2 predominantly constitute cluster 2. In contrast, the majority of other individuals exhibit mixed genomic profiles with no clear assignment to either group. These findings underscore a generally poor genetic structuring across the dataset, with pervasive admixture that limits the resolution of distinct clusters.

Genomic analyses detected moderate to relatively high levels of genetic diversity for the studied populations (Table 5), ranging from the highest levels of Mendizar-P2 (Ho=0.219, Hs=0.277) and Errozate-I2 (Ho=0.213, Hs=0.276) and the lowest of Errozate-P1 (Ho=0.179, Hs=0.270) and Urkulu-P3 (Ho=0.192, Hs=0.278) but with small differences. Genetic diversity was highest for superdominant (Ho=0.208, Hs=0.275) and lowest for stand populations (Ho=0.196, Hs=0.275) but showed small variations in all cases. The pairwise Fst values (Table S5) revealed relatively low genomic differentiation among populations, with values ranging from 0.124 (Ezkanda-S1/Arpea-I1) to 0.193 (Orion-S2/Mendizar-I3) for individual populations (Table S5). FIS values were consistently and significantly positive across populations, ranging from 0.211 to 0.34 with an overall mean of 0.269, which indicates a heterozygote deficit relative to Hardy–Weinberg expectations. In parallel, the low selfing rates (overall sC=C0.417) reinforce the predominantly outcrossing mating system observed in this species (Table 5). At the expansion phase level, differences in mating dynamics were also evident. Specifically, superdominant populations displayed a lower selfing rate (sC=C0.387) and a reduced FIS value (0.242) compared to the scattered populations (sC=C0.431; FISC=C0.282) and stand populations (sC=C0.440; FISC=C0.289). Both standard and hierarchical AMOVA analyses showed that the highest proportion of diversity was found within populations (around 96%), while the proportion of diversity and differentiation between the hierarchical groups tested were not significant (Table 6).

**Table 5.** Estimates of genetic diversity of *B. rupestre* analysed by populations, and mean values for geographic areas and expansion phases. Ho, observed heterozygosity; Hs, expected heterozygosity; s, selfing rate; FIS, inbreeding coefficient and associated P-value.

| SITES | Ho | Hs | s | FIS | P-value |
| --- | --- | --- | --- | --- | --- |
| Ezkanda-S1 | 0.212 | 0.277 | 0.372 | 0.235 | < 0.001 |
| Orion-S2 | 0.194 | 0.275 | 0.45 | 0.295 | < 0.001 |
| Urkulu-S3 | 0.192 | 0.275 | 0.454 | 0.302 | < 0.001 |
| Errozate-P1 | 0.179 | 0.27 | 0.502 | 0.34 | < 0.001 |
| Mendiziar-P2 | 0.219 | 0.277 | 0.338 | 0.211 | < 0.001 |
| Urkulu-P3 | 0.191 | 0.278 | 0.474 | 0.311 | < 0.001 |
| Arpea-I1 | 0.212 | 0.277 | 0.373 | 0.233 | < 0.001 |
| Errozate-I2 | 0.213 | 0.276 | 0.368 | 0.228 | < 0.001 |
| Mendiziar-I3 | 0.2 | 0.272 | 0.418 | 0.265 | < 0.001 |
| GEOGRAPHICAL AREA | Ho | Hs | s | FIS | P-value |
| Arpea | 0.212 | 0.277 | 0.373 | 0.233 | < 0.001 |
| Errozate | 0.197 | 0.273 | 0.429 | 0.279 | < 0.001 |
| Ezkanda | 0.212 | 0.277 | 0.372 | 0.235 | < 0.001 |
| Mendiziar | 0.208 | 0.274 | 0.383 | 0.242 | < 0.001 |
| Orion | 0.194 | 0.275 | 0.45 | 0.295 | < 0.001 |
| Urkulu | 0.192 | 0.276 | 0.463 | 0.306 | < 0.001 |
| EXPANSION PHASE | Ho | Hs | s | FIS | P-value |
| Scattered (S) | 0.198 | 0.275 | 0.431 | 0.282 | < 0.001 |
| Stand (P) | 0.196 | 0.275 | 0.44 | 0.289 | < 0.001 |
| Superdominant (I) | 0.208 | 0.275 | 0.387 | 0.242 | < 0.001 |
| OVERALL | Ho | Hs | s | FIS | P-value |
| *B. rupestre* | 0.201 | 0.275 | 0.417 | 0.269 | < 0.001 |

**Table 6.** Analysis of the molecular variance (AMOVA) of *B. rupestre* populations. Hierarchical AMOVAs established according to populations, geographical areas and expansion phases.

STANDARD AMOVA (SITES)
| Source of variation | d.f. | Percentage of variation | P-value |
| --- | --- | --- | --- |
| Among sites | 8 | 3.47 | <0.001 |
| Within sites | 90 | 96.53 | 1 |
| Total | 98 | 100 |  |

HIERARCHICAL AMOVAs GEOGRAPHICAL AREA
| Source of variation | d.f. | Percentage of variation | P-value |
| --- | --- | --- | --- |
| Among zones | 5 | 2.45 | <0.001 |
| Between sites within zones | 84 | 0.00 | 1 |
| Within sites | 90 | 97.55 | 1 |
| Total | 179 | 100 |  |

EXPANSION PHASE
| Source of variation | d.f. | Percentage of variation | P-value |
| --- | --- | --- | --- |
| Among expansion phases | 2 | 1.04 | <0.001 |
| Between sites within expansion phase | 87 | 0 | 1 |
| Within sites | 90 | 98.96 | 1 |
| Total | 179 | 100 |  |

MANAGEMENT REGIMES
| Source of variation | d.f. | Percentage of variation | P-value |
| --- | --- | --- | --- |
| Among management regimes | 1 | 0.62 | <0.001 |
| Between sites within management regimes | 88 | 0 | 1 |
| Within sites | 90 | 99.38 | 1 |
| Total | 179 | 100 |  |

### 3.4 Genetic clonality

A total of 67 MLGs (and 67 MLLs) were detected among the 90 sampled individuals of *B. rupestre* (Table 7, Figure S2). The genetic threshold distance, under which two MLGs were considered to belong to the same MLL, was estimated using the farthest neighbour method, which was found to be the optimal estimator (Figure S3). The POPPR analysis found a different number of clones within populations, geographic areas and expansion phases. Between six and nine MLLs were detected in the analysed populations (Table 7). The Errozate-P1 and Orion-S2 populations showed the lowest Shannon-Weiner diversity index (H=1.7) and clonal richness (R=0.556), along with the Ezkanda-S1 and Urkulu-S3 populations (H=1.75-1.89; R=0.750-0.833), while Mendizar-P2 and Urkulu-P3 had the highest (H=2.04; R=0.875), followed by the superdominant populations Arpea-I1, Errozate-I2 and Mendizar-I3 (H=2.15-2.02; R=0.800-0.636) (Table 7). When genets within the population propagated preferentially clonally, lower MLG clonal richness resulted. Consequently, Errozate-P1 and Orion-S2 were the populations with the most clonal individuals per MLG. The latter populations plus Urkulu-S3 had the highest standardized association index values, and all populations showed r_d_ values that significantly deviated from free recombination.

**Table 7.** Clonality descriptors for the studied populations, including number of individuals (N), multilocus lineages (MLL), expected multilocus lineages (eMLL), Shannon-Weiner index (H), Simpson index corrected for population size (Lambda), clonal Evenness index (E.5), standardized association index (rd), and clone richness (R). The values were analysed by populations, as well as by geographic areas and expansion phases. Asterisks in the association index show the significance level (p<0.005).

| SITES | N | MLL | eMLL | H | Lambda | E.5 | rd | R |
| --- | --- | --- | --- | --- | --- | --- | --- | --- |
| Ezkanda-S1 | 7 | 6 | 6 | 1.75 | 0.952 | 0.94 | 0.021* | 0.833 |
| Orion-S2 | 10 | 6 | 6 | 1.70 | 0.889 | 0.90 | 0.037* | 0.556 |
| Urkulu-S3 | 10 | 7 | 7 | 1.83 | 0.911 | 0.87 | 0.044* | 0.667 |
| Errozate-P1 | 10 | 6 | 6 | 1.70 | 0.889 | 0.90 | 0.039* | 0.556 |
| Mendiziar-P2 | 9 | 8 | 8 | 2.04 | 0.972 | 0.95 | 0.012* | 0.875 |
| Urkulu-P3 | 9 | 8 | 8 | 2.04 | 0.972 | 0.95 | 0.018* | 0.875 |
| Arpea-I1 | 11 | 9 | 8.36 | 2.15 | 0.964 | 0.94 | 0.011* | 0.800 |
| Errozate-I2 | 12 | 9 | 7.95 | 2.14 | 0.955 | 0.94 | 0.012* | 0.727 |
| Mendiziar-I3 | 12 | 8 | 7.27 | 2.02 | 0.939 | 0.95 | 0.017* | 0.636 |
| GEOGRAPHICAL AREA | N | MLL | eMLL | H | Lambda | E.5 | rd | R |
| Arpea | 11 | 9 | 8.36 | 2.15 | 0.964 | 0.94 | 0.010* | 0.800 |
| Errozate | 22 | 15 | 8.52 | 2.63 | 0.965 | 0.92 | 0.008* | 0.667 |
| Ezkanda | 7 | 6 | 6 | 1.75 | 0.952 | 0.94 | 0.021* | 0.833 |
| Mendiziar | 21 | 16 | 8.93 | 2.71 | 0.976 | 0.94 | 0.006* | 0.750 |
| Orion | 10 | 6 | 6 | 1.70 | 0.889 | 0.9 | 0.038* | 0.556 |
| Urkulu | 19 | 15 | 8.81 | 2.63 | 0.971 | 0.89 | 0.016* | 0.778 |
| EXPANSION PHASE | N | MLL | eMLL | H | Lambda | E.5 | rd | R |
| Scattered (S) | 27 | 19 | 19 | 2.85 | 0.972 | 0.89 | 0.010* | 0.692 |
| Stand (P) | 28 | 22 | 21.4 | 3.02 | 0.981 | 0.91 | 0.006* | 0.778 |
| Superdominant (I) | 35 | 26 | 21.7 | 3.20 | 0.985 | 0.94 | 0.003* | 0.735 |
| OVERALL | N | MLL | eMLL | H | Lambda | E.5 | rd | R |
| *B. rupestre* | 90 | 67 | 10 | 4.13 | 0.994 | 0.93 | 0.001* | 0.750 |

## 4 Discussion

### 4.1 Investment of *B. rupestre* in sexual reproduction under different management regimes associated to different expansion phases

The impact of herbivores on the sexual reproduction of grasses has been a subject of debate in scientific literature. A recent review of Wentao et al. (2023) suggests that herbivory promotes clonal propagation through tillering, at the expense of sexual reproduction. We found the highest standardized association index (r_d_) in populations at grazed sites and also fewer inflorescences and a lower number of seeds than in the rest of sites (3.0 *vs.* 10.6-12.7 inflorescences/m^2^, and 2.8 *vs.* 19.0-24.2 seeds/m^2^). The timing of defoliation is key to understanding the mechanisms of inhibition of sexual reproduction and reproductive development of tillers in grasses. When defoliation occurs at the beginning of stem elongation, the apical meristem becomes accessible to the herbivore’s mouth, potentially leading to the loss of the developing inflorescence (Hopkins et al., 2003). Less inflorescences and seeds per spikelet were collected from grazed locations than from the other sites. Low seed yields related to herbivory have been reported in former reviews (Anderson & Frank, 2003; Wentao et al., 2023) and may indicate low fecundity and failed seed development, as well as physical removal of seeds from the inflorescence by grazers. Repeated clipping experiments on *Bromus* spp. reported a significant negative effect on seed yields (Hempy-Mayer & Pyke, 2008). Regarding seed removal, herbivores can greatly contribute to seed dispersal by ingesting seeds (endozoochory) or transporting them through their fur and hooves (epizoochory) (Albert et al., 2015; Couvreur et al., 2004; Milotić et al., 2017). In this regard, the role of scattered shrubs in grazed areas in protecting the sexual reproductive structures of grasses warrants further exploration. Shrubs with deterrent structures, such as the thorny *Ulex gallii* in the study area, may act as shelters for grasses growing within, providing hot spots of sexual offsprings for the surrounding grazing environment (biogenic refuges; Milchunas & NoyCMeir, 2002; Rebollo et al., 2002).

Fire is thought to have a positive impact on the sexual reproduction of grasses, including low-intensity prescribed burns (Dyer, 2002). *B. retusum,* a perennial species closely related to *B. rupestre* that thrives in fire-prone Mediterranean environments (Eugenio & Lloret, 2006; Santana et al., 2013), significantly increases inflorescence production and seed production after fires, particularly summer fires (Vidaller et al., 2019; Vilà-Cabrera et al., 2008). In the study area, where burnings are applied in winter and buds and rhizomes are protected from flames, it is unclear whether fire will have a positive effect on flowering (Canals et al., 2014; Clarke et al., 2013). Our seed yields did not differ significantly between superdominant (recurrently burned) and stand populations, with the latter even developing a higher number of inflorescences. Stand populations of *B. rupestre* are avoided by herbivores, creating plant refuges that allow individuals to successfully complete their reproductive cycle. In other perennial grasses constituting dense stands, like *Festuca idahoensis*, sexual reproduction has been found to be surprisingly more important than clonal propagation for stand expansion (Liston et al., 2003), a result that may certainly contribute to maintaining high levels of genetic diversity in the stands.

### 4.2 Genomic diversity and structure of the *B. rupestre* populations

Our genomic study has uncovered both genomic divergence of the nine surveyed populations of *B. rupestre* and a trend of genomic homogeneity among them (Figures 5, 6; Tables 5, S5). The clear phylogenomic divergence of the populations, which showed a great match with their west-central/south-east spatial geographical distribution (Figure 5), is a consequence of the high resolution of the ddRAD SNP data (Peterson et al., 2012). SNPs provide a reliable genome-wide picture, recovering subtle patterns of micro-scale phylogeographic differentiation and also informing local population structure (Dufresnes et al., 2023). Despite this, the genomic structure data indicated weak structuring for the *B. rupestre* populations, which was also consistent with the polyphyly of some of the Orion-S2, Ezkanda-S1 and Errozate-P1 individuals (Figure 6). This tendency towards genomic homogeneity among populations, caused by gene flow, fits the expectations of this strongly outcrossing and wind-pollinated plant (sub *B. pinnatum* 2n=28; Catalan et al., 2016) in the open environments of the Aezkoa mountains where there are no geographic barriers that separate spatially close populations. Landscape genetic models have demonstrated that wind shapes large-scale genetic patterns of wind-pollinated plants, with populations linked by stronger winds being more genetically similar (Kling & Ackerly, 2021). Studies of cross-pollination distances of grasses also indicated that gene flow is mainly affected by the distance between donor and acceptor plants and the density of acceptor plants (Rognli et al., 2000). Landscape genetic analyses of other wind-pollinated Pyrenean grasses (e.g. *Festuca eskia*) also found largely homogeneous genetic groups, encompassing distant populations separated by more than 25 km linear distance (Catalán et al., 2013). Similarly, a general lack of isolation by distance and weak genetic population structuring was found for the close congener *B. pinnatum* in 100 km^2^ of the Jura mountains highlands (Bąba et al., 2012). This fact reflects a common pattern of genomic homogenization and rapid expansion of invasive species of the *B. pinnatum-B. rupestre* complex. The best *K*=2 genetic groups detected within the *B. rupestre* samples studied indicated a genetic mixture and admixture of genetic profiles in most populations. The predominance of one or another genetic group does not correspond to spatially close populations, suggesting that directionally unbalanced winds may have caused asymmetric proportions of gene flow (Kling & Ackerly, 2021), or that local topography or other features could have influenced effective pollination and dispersal rates (Cruzan & Hendrickson, 2020). The absence of strong genetic structure within the studied *B. rupestre* populations was also reflected in AMOVAs that did not detect significant structuring for any hierarchical clustering related to the geographic area or expansion phases (Table 6). These results suggest that the current human-mediated practices likely do not counteract the predominant effect of genetically homogenizing wind pollination and potential selective pressures underlying local adaptation patterns (Dauphin et al., 2023).

In contrast, the levels of genomic diversity of *B. rupestre* populations were generally relatively high compared to those of other plant species also studied with RADseq SNP markers (Dong et al., 2022). The Ho and Hs values ranged from moderate (Errozate-P1) to relatively high (Mendizar-P2) (Table 5), as expected for this preferentially allogamous allotetraploid species (Catalan et al., 2016). Consequently, the inbreeding coefficient and selfing rates were very low, which is also in line with the predominant outbreeding reproductive system of this species that shows extremely low self-pollination rates (Khan & Stace, 1999). The observed and expected heterozygosity values of the *B. rupestre* populations are also higher than those estimated for its annual selfing Mediterranean congeners, the diploids *B. distachyon* (Marques et al., 2017, SSR data) and *B. stacei* (Campos et al., 2024, RADseq data), and the allotetraploid *B. hybridum* (Shiposha et al., 2020, SSR data), and similar to or slightly higher than those of the selfing Eurasian perennial diploid congener *B. sylvaticum* (Rosenthal et al., 2008), according to these studies. The genomic diversity of the *B. rupestre* populations could be considered equivalent to that of their close allogamous congener *B. pinnatum*. However, the genetic diversity estimator for *B. pinnatum*, calculated from AFLP markers, showed extremely high values (Bąba et al., 2012). It is important to note that diversity estimates derived from AFLP presence/absence data may not be directly comparable to those obtained from SNPs’ allelic frequencies due to differences in the underlying methodologies and algorithms used for these markers. The relative high levels of genomic diversity detected in Aezkoan populations of *B. rupestre* do not seem to have been affected by the mixed presence of genets and ramets (MLG clones) in the same locations (Table 7), although the coexistence of asexuals does not affect the reproductive success of the genets, as evidenced by high seed production rates. FIS analysis indicates that Aezkoan *B. rupestre* populations deviate significantly from HWE, suggesting a predominance of non-random mating, potentially due to partial selfing or subtle population substructure, highlighting that gene flow may not be sufficient to fully counteract the effects of reproductive system constraints or local isolation.

### 4.3 Genetic diversity and clone richness of *B. rupestre* under different management regimes are associated to different expansion phases

In this study, the majority of genetic diversity was found within populations, and no significant genetic differentiation was detected among expansion phases associated with different management practices (herbivory, abandonment, and burning). However, several patterns emerged for discussion. Stand, scattered and superdominant populations displayed overall high genetic diversity, characterized by high heterozygosity and low inbreeding coefficients (FIS) (Table 5). In contrast, the highest number of distinct asexual clones (26 multilocus lineages [MLLs] compared to the expected 21.7) was observed in superdominant populations, while stand and scattered populations had significantly lower number of MLLs (Table 7). The results obtained may suggest a *bias* due to the sampling scale. Since there are many more individuals of *B. rupestre* in populations in intermediate and advanced stages of invasion, the probability of identifying a high number of clones increases. Interestingly, the superdominant populations with significantly higher number of asexual clones also showed slightly higher though non-significant sexually derived heterozygosity values (Tables 5, 7). The weak relationship between heterozygosity and the number of multilocus lineages in superdominant populations, may suggest a complex pattern of clonality variability influenced by historical environmental factors shaping each population (e.g., burns, herbivory regimes), and their co-existence with sexually different genotypes.

For scattered populations, herbivore-driven seed dispersal and improved seedling establishment opportunities can enhance genetic diversity, even with the dominance of vegetative expansion and a low proportion of sexual structures. Research has shown that moderate grazing positively impacts seedling recruitment and offspring diversity (Gao-Lin et al., 2011). Although herbivory reduced *B. rupestre* seed propagules in scattered populations, it might facilitate seedling recruitment by reducing competition and creating spaces for seedling establishment and germination (Gao-Lin et al., 2011; Gartner et al., 1983; Kiviniemi & Eriksson, 1999).

In superdominant populations, a significant investment in sexual reproductive organs was observed despite the presence of rhizomes and buds that facilitate vegetative reproduction. Comparable patterns have been documented in closely related species, such as *B. pinnatum*, which exhibit successful vegetative (Baba et al., 2012; Bobbink & Willems, 1987) and sexual reproduction (Schlaepfer, 1997; Schläpfer & Fischer, 1998), and in *B. sylvaticum* (Novak & Welfley, 1997; Rosenthal et al., 2008). Clonal growth enables adaptation to environmental heterogeneity, alleviates local resource scarcity, and buffers against environmental stresses and disturbances. However, clonality carries long-term costs, including the inability to generate genetic variation in offspring, increased risk of mutation accumulation due to a lack of recombination, and limited propagule dispersal to new colonizing sites (Lamont & Wiens, 2003; Nathan & Muller-Landau, 2000). As a result, the combination of sexual and asexual reproduction, leading to a high number of distinct genotypes and clones, seems crucial for the success of these species, as reported for other invasive species (Barrett et al., 2008; Van Kleunen et al., 2010).

## 5 Conclusions

The results of this study show a high genetic diversity of *B. rupestre* populations, with asexual reproduction in all the populations, coupled with sexual mating, and a weak genomic structure due to the absence of barriers to gene flow among high-altitude populations. The hypothesis that populations at different expansion phases and exposed to different disturbance regimes would develop distinct genomic identities is not supported by the results. Rather, the genetic differentiation and divergence of the populations found are a consequence of life history and spatial isolation, even within the restricted geographic area of the Aezkoa valley. The weak relationship between genetic diversity (heterozygosity) and the number of different clones among populations, suggests a complex pattern of genotypic and clonal variability, which may be influenced by environmental factors (historical disturbance regimes) affecting each population.

## Acknowledgments

We thank the Aezkoa Valley Council for permissions for sampling in the study sites, L. Múgica, M. Saldise and C. Segura for support in the field work, and Luis A Inda and Ma Angeles Decena for their help with flow cytometry genome size analysis. ddRADseq data from the studied *B. rupestre* were generated at Floragenex, INC. The population genomics and evolutionary analyses were performed in the Bioflora laboratory of the Escuela Politécnica Superior de Huesca (EPSHU, Universidad de Zaragoza, Spain).

## Author Contributions

The research was conceptualized by RM, LS, PC and MD. MD and LS designed the experimental set-up and MD and RM did the sampling monitoring in the Aezkoa valley. The genetic data were processed and analyzed by MC, PC, MD, SB, AS and the reproductive data by MD and LS. The writing process was leaded by MD, RM, MC and PC and all authors reviewed the manuscript. Supervising, project administration and funding acquisition depended on RM, PC and LS.

## Data Availability Statement

The input and output data generated for this study can be found at Github [https://github.com/Bioflora/Brupestre_ddRAD]

