## Supplementary figures and images for "Exploring genomic diversity and reproductive strategies in three expansion phases of the superdominant *Brachypodium rupestre* in high mountain grasslands"

### Figure S1

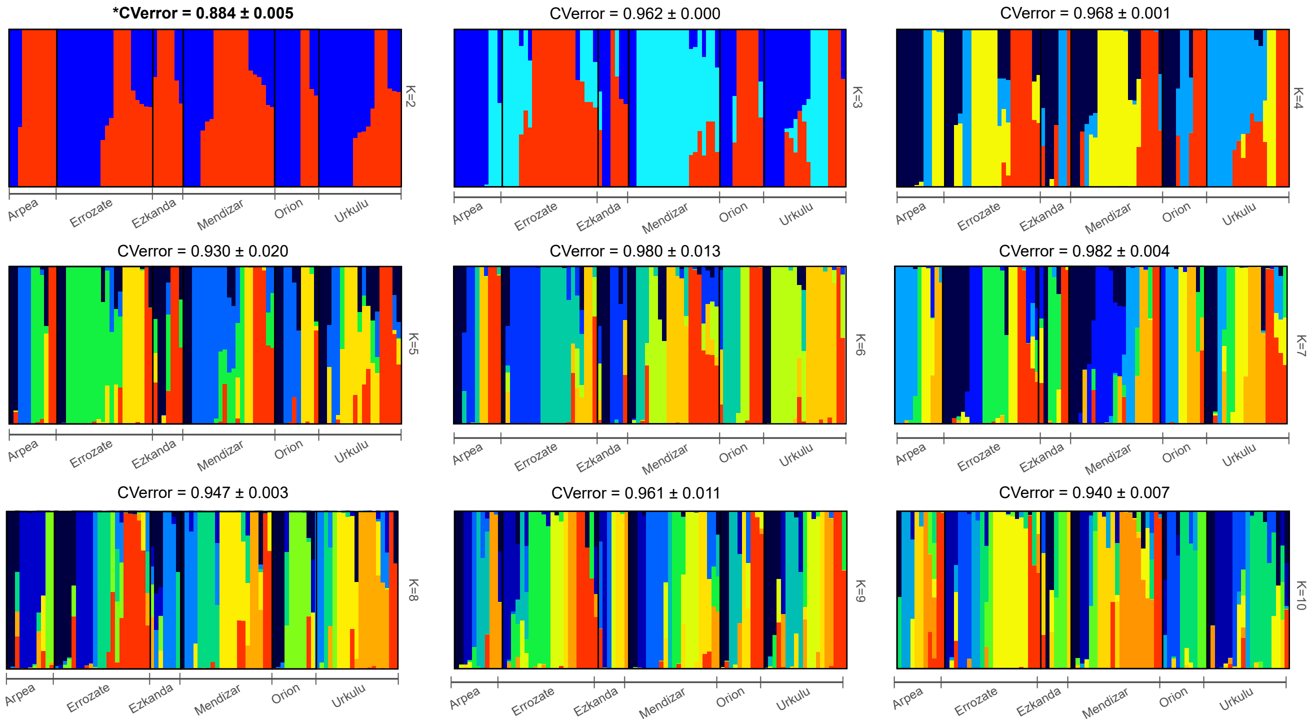

### Figure S2

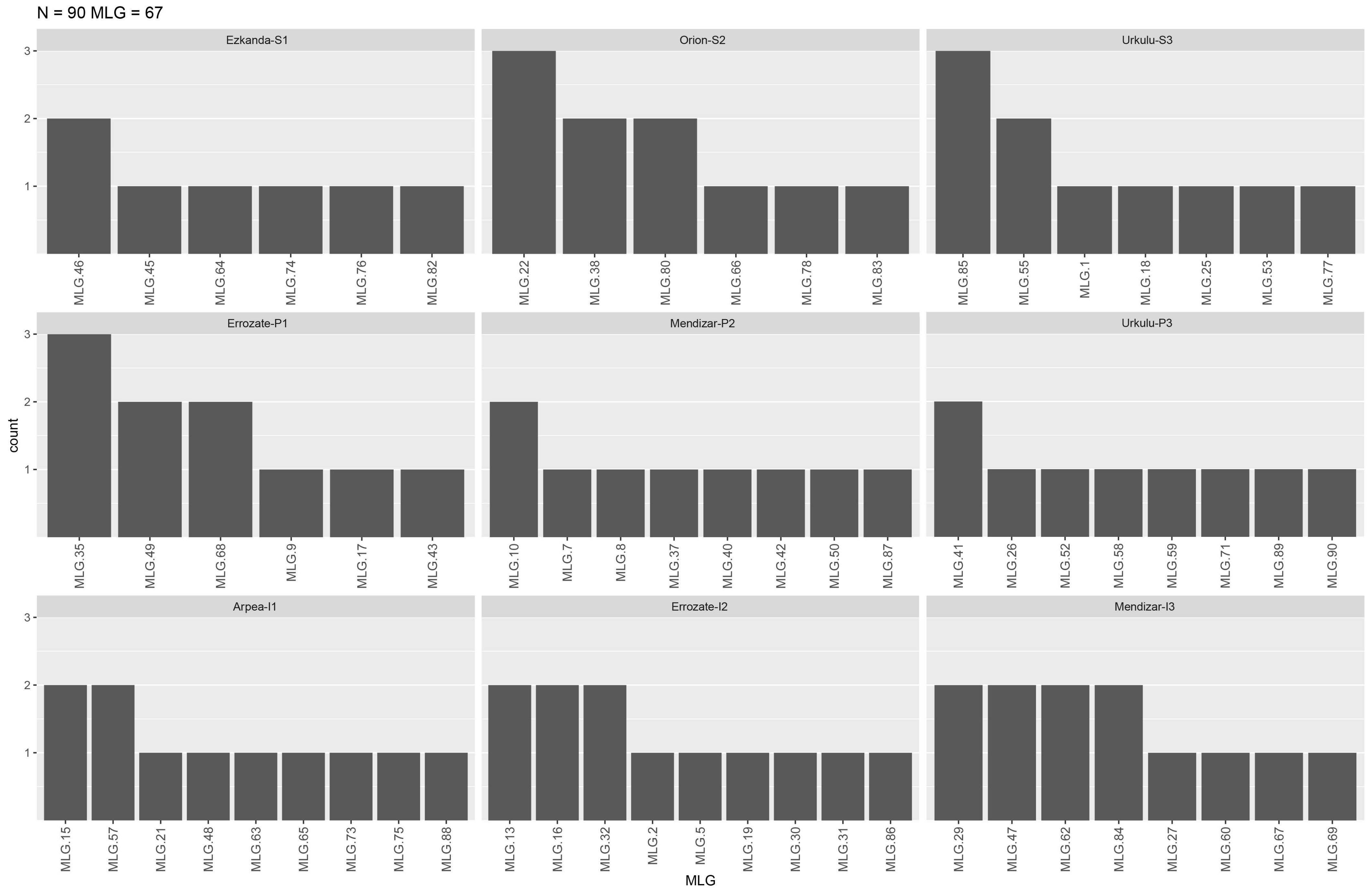

### Figure S3

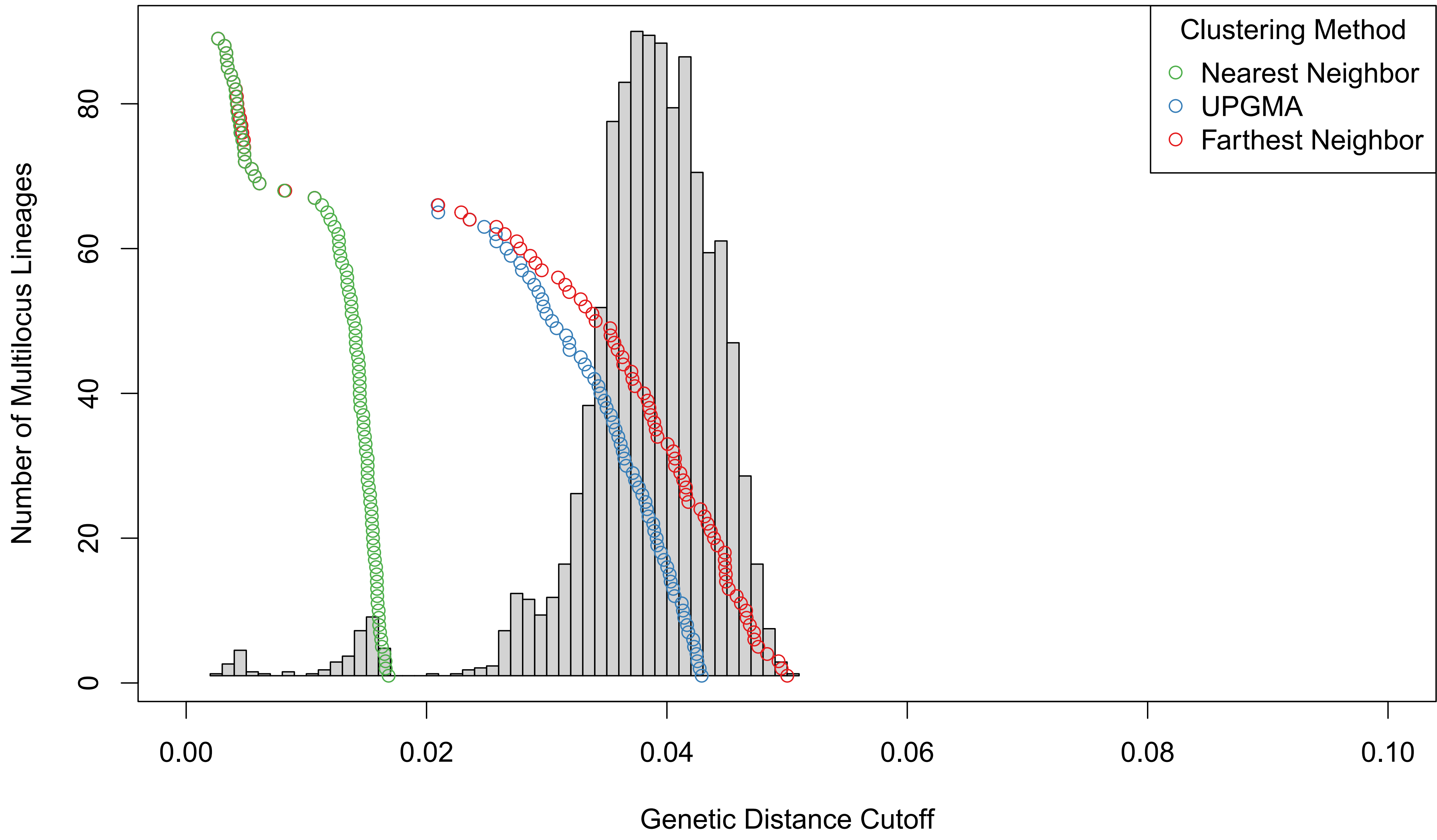
