## Supplementary material for "Exploring genomic diversity and reproductive strategies in three expansion phases of the superdominant *Brachypodium rupestre* in high mountain grasslands": Table S1

**Supplementary Table S1**. Coordinates of the sampling sites.

|  |  |  |  | Coordinates (UTM Zone 30N) | |
| --- | --- | --- | --- | --- | --- |
|  | Site | Expansion phase |  | X | Y |
| 1 | Ezkanda-S1 | Scattered |  | 346802.669 | 4765098.531 |
| 2 | Orion-S2 |  |  | 347415.412 | 4763073.788 |
| 3 | Urkulu-S3 |  |  | 342449.461 | 4767798.543 |
| 4 | Errozate-P1 | Stand |  | 348545.360 | 4766705.215 |
| 5 | Mendizar-P2 |  |  | 350618.286 | 4764183.069 |
| 6 | Urkulu-P3 |  |  | 342680.827 | 4768184.324 |
| 7 | Arpea-I1 | Invading |  | 347459.750 | 4766413.805 |
| 8 | Errozate-I2 |  |  | 348383.334 | 4766226.623 |
| 9 | Mendizar-I3 |  |  | 350899.785 | 4764791.340 |
