## Supplementary material for "Exploring genomic diversity and reproductive strategies in three expansion phases of the superdominant *Brachypodium rupestre* in high mountain grasslands": Table S2

**Table S2** Population genomic study of *Brachypodium rupestre* in the Aezkoa Valley (Navarra, Spain) using ddRADseq data. Raw reads used in Ipyrad (Raw), reads that passed filters (Filtered), clusters created from those filtered reads (Clusters), reads used for consensus (Consensus), and final loci assembled per sample (Assembled).

| **Sample** | **Raw** | **Filtered** | **Clusters** | **Consensus** | **Assembled** |
| --- | --- | --- | --- | --- | --- |
| Ezkanda-S1_1 | 18259 | 18236 | - | - | - |
| Ezkanda-S1_2 | 5471337 | 5468189 | 168117 | 71637 | 48035 |
| Ezkanda-S1_3 | 5345400 | 5342193 | 199239 | 78357 | 52313 |
| Ezkanda-S1_4 | 5135197 | 5132055 | 199547 | 75086 | 50480 |
| Ezkanda-S1_5 | 4637280 | 4634415 | 206610 | 77669 | 51666 |
| Ezkanda-S1_6 | 5634704 | 5631271 | 177331 | 73714 | 49400 |
| Ezkanda-S1_7 | 83818 | 83735 | - | - | - |
| Ezkanda-S1_8 | 5512441 | 5509262 | 190891 | 80577 | 54639 |
| Ezkanda-S1_9 | 6257241 | 6253714 | 200787 | 88123 | 61451 |
| Orion-S2-10 | 4737450 | 4734253 | 214362 | 81765 | 55179 |
| Orion-S2-11 | 4096816 | 4094028 | 194228 | 69867 | 46517 |
| Orion-S2-12 | 4525423 | 4522404 | 187212 | 73870 | 49542 |
| Orion-S2-13 | 5770811 | 5767047 | 223854 | 81537 | 54544 |
| Orion-S2-14 | 4846873 | 4843677 | 197183 | 79092 | 53361 |
| Orion-S2-15 | 4859753 | 4856512 | 218352 | 77651 | 52052 |
| Orion-S2-16 | 5074441 | 5071311 | 188764 | 77527 | 52007 |
| Orion-S2-17 | 5229566 | 5226375 | 213273 | 77097 | 51279 |
| Orion-S2-18 | 5378739 | 5375309 | 205161 | 83033 | 56510 |
| Orion-S2-19 | 5040663 | 5037362 | 225314 | 81557 | 54856 |
| Urkulu-S3_20 | 5337174 | 5333606 | 210636 | 86958 | 58882 |
| Urkulu-S3_21 | 3649887 | 3647488 | 165208 | 64851 | 42658 |
| Urkulu-S3_22 | 4991024 | 4987856 | 209298 | 82044 | 55482 |
| Urkulu-S3_23 | 5373803 | 5370480 | 203116 | 87506 | 59528 |
| Urkulu-S3_24 | 1771468 | 1769880 | 119919 | 32772 | 15561 |
| Urkulu-S3_25 | 4621220 | 4618304 | 206359 | 72603 | 48494 |
| Urkulu-S3_26 | 5151273 | 5148019 | 200777 | 76384 | 51171 |
| Urkulu-S3_27 | 5044945 | 5041705 | 204311 | 77658 | 52368 |
| Urkulu-S3_28 | 5855878 | 5852117 | 198126 | 81699 | 55393 |
| Urkulu-S3_29 | 5137329 | 5133991 | 203807 | 80205 | 54023 |
| Errozate-P1_30 | 4621430 | 4618382 | 200653 | 77148 | 52047 |
| Errozate-P1_31 | 4712994 | 4709846 | 189229 | 78201 | 53134 |
| Errozate-P1_32 | 4690808 | 4687817 | 192287 | 76313 | 51446 |
| Errozate-P1_33 | 4796698 | 4793473 | 219651 | 78022 | 52802 |
| Errozate-P1_34 | 4293403 | 4290440 | 276653 | 75954 | 49437 |
| Errozate-P1_35 | 4373246 | 4370395 | 199741 | 75521 | 51101 |
| Errozate-P1_36 | 3142265 | 3140119 | 174878 | 61372 | 40252 |
| Errozate-P1_37 | 4324710 | 4321818 | 204762 | 77249 | 52139 |
| Errozate-P1_38 | 4083727 | 4080946 | 216493 | 75157 | 50425 |
| Errozate-P1_39 | 5201960 | 5198727 | 225863 | 80994 | 54638 |
| Mendizar-P2_40 | 4847354 | 4844246 | 275123 | 78414 | 52058 |
| Mendizar-P2_41 | 5525385 | 5521746 | 223580 | 89126 | 60629 |
| Mendizar-P2_42 | 5504005 | 5500463 | 238283 | 87122 | 59133 |
| Mendizar-P2_43 | 5492666 | 5489264 | 227185 | 90005 | 62409 |
| Mendizar-P2_44 | 4536298 | 4533527 | 208515 | 85408 | 58863 |
| Mendizar-P2_45 | 213200 | 213070 | - | - | - |
| Mendizar-P2_46 | 5209577 | 5206129 | 188763 | 82970 | 56076 |
| Mendizar-P2_47 | 5620098 | 5616784 | 186374 | 85712 | 58755 |
| Mendizar-P2_48 | 4788553 | 4785459 | 198043 | 84419 | 57998 |
| Mendizar-P2_49 | 4732250 | 4729111 | 279997 | 78916 | 51723 |
| Urkulu-P3_50 | 5741719 | 5738398 | 185524 | 83219 | 56478 |
| Urkulu-P3_51 | 5920756 | 5917338 | 199960 | 87017 | 59156 |
| Urkulu-P3_52 | 6229160 | 6225631 | 174359 | 75910 | 51731 |
| Urkulu-P3_53 | 5796538 | 5793304 | 175689 | 77806 | 53384 |
| Urkulu-P3_54 | 4972660 | 4969995 | 175909 | 75577 | 51862 |
| Urkulu-P3_55 | 3146759 | 3144845 | 145963 | 54096 | 34740 |
| Urkulu-P3_56 | 6852 | 6849 | - | - | - |
| Urkulu-P3_57 | 6133451 | 6130231 | 172221 | 73939 | 49614 |
| Urkulu-P3_58 | 5394979 | 5391817 | 167419 | 67993 | 45218 |
| Urkulu-P3_59 | 6013270 | 6009964 | 177847 | 73677 | 49356 |
| Arpea-I1_60 | 5338644 | 5335538 | 188805 | 72812 | 48483 |
| Arpea-I1_61 | 5094189 | 5091042 | 223805 | 76491 | 51251 |
| Arpea-I1_62 | 5608643 | 5605432 | 192698 | 75520 | 50564 |
| Arpea-I1_63 | 6270790 | 6267236 | 183297 | 81560 | 55100 |
| Arpea-I1_64 | 5586280 | 5583175 | 179648 | 81766 | 55804 |
| Arpea-I1_65 | 4941280 | 4938364 | 176045 | 73810 | 49420 |
| Arpea-I1_66 | 5748637 | 5745422 | 208090 | 82724 | 57125 |
| Arpea-I1_67 | 6251137 | 6247529 | 182695 | 79924 | 54267 |
| Arpea-I1_68 | 5253513 | 5250493 | 204151 | 80889 | 55260 |
| Arpea-I1_69 | 6325572 | 6321897 | 168385 | 76989 | 52278 |
| Arpea-I1_70 | 7557 | 7547 | - | - | - |
| Arpea-I1_71 | 5064676 | 5061641 | 168196 | 69783 | 46884 |
| Errozate-I2_72 | 5441479 | 5437924 | 220980 | 86928 | 59867 |
| Errozate-I2_73 | 4889991 | 4886865 | 195304 | 80369 | 54891 |
| Errozate-I2_74 | 5502843 | 5499408 | 201016 | 87201 | 59680 |
| Errozate-I2_75 | 5677934 | 5674436 | 208703 | 86599 | 59084 |
| Errozate-I2_76 | 5496423 | 5492997 | 194764 | 81755 | 55391 |
| Errozate-I2_77 | 5492168 | 5488556 | 218582 | 90461 | 62588 |
| Errozate-I2_78 | 5062135 | 5058870 | 198665 | 82417 | 56280 |
| Errozate-I2_79 | 5004466 | 5001261 | 194397 | 79937 | 54387 |
| Errozate-I2_80 | 4962071 | 4958849 | 197820 | 80578 | 54425 |
| Errozate-I2_81 | 4872151 | 4868947 | 211636 | 86079 | 57241 |
| Errozate-I2_82 | 4058310 | 4055580 | 197648 | 70563 | 46848 |
| Errozate-I2_83 | 5116421 | 5113090 | 208783 | 80517 | 54640 |
| Mendizar-I3_84 | 5183577 | 5180210 | 194965 | 78638 | 53385 |
| Mendizar-I3_85 | 5032846 | 5029519 | 225886 | 79047 | 53474 |
| Mendizar-I3_86 | 4450184 | 4447346 | 192238 | 71644 | 47454 |
| Mendizar-I3_87 | 4108271 | 4105576 | 195653 | 74042 | 48787 |
| Mendizar-I3_88 | 4714299 | 4711290 | 226439 | 79709 | 53719 |
| Mendizar-I3_89 | 3271206 | 3269000 | 195742 | 62006 | 38948 |
| Mendizar-I3_90 | 4180362 | 4177517 | 204810 | 72189 | 46941 |
| Mendizar-I3_91 | 3897705 | 3895021 | 199403 | 71467 | 44947 |
| Mendizar-I3_92 | 3388622 | 3386359 | 172544 | 65459 | 43043 |
| Mendizar-I3_93 | 4204221 | 4201411 | 214596 | 75640 | 49515 |
| Mendizar-I3_94 | 3013845 | 3011753 | 166687 | 61492 | 39834 |
| Mendizar-I3_95 | 4447907 | 4444805 | 246253 | 75070 | 49464 |
