## Supplementary material for "Exploring genomic diversity and reproductive strategies in three expansion phases of the superdominant *Brachypodium rupestre* in high mountain grasslands": Table S3

**Table S3**. Genome size analysis. Flow cytometry results for nuclear DNA content estimation (pg/2C) across six individual samples of Brachypodium rupestre collected from the Arpea and Urkulu populations of the Aezkoa valley (Navarra, Spain). Measurements were performed in two replicates (1 and 2) for each sample (Brup1 to Brup6) using Solanum lycopersicum L. “Stupické polní rané” (1.96 pg/2C) as internal standard. The table indicates the total number of nuclei analyzed (nucleids), the mean fluorescence value (mean), the coefficients of variation (CV), and the estimated genome size (pg/2C). The final DNA content (pg/2C) was averaged for each population.

| **Population** | **Samples** | **Nucleids** | **Mean** | **CV** | **pg/2C** | **pg/2C Average** |
| --- | --- | --- | --- | --- | --- | --- |
| Arpea | Brup1_1 | 1422 | 17236.26 ± 398.16 | 2.31 | 1.37 | 1.410 ± 0.029 |
|  | Brup1_2 | 4241 | 16817.65 ± 484.35 | 2.88 | 1.39 |  |
|  | Brup2_1 | 1597 | 17887.42 ± 418.57 | 2.34 | 1.41 |  |
|  | Brup2_2 | 1996 | 16363.41 ± 456.54 | 2.79 | 1.43 |  |
|  | Brup3_1 | 1620 | 17132.05 ± 416.31 | 2.43 | 1.41 |  |
|  | Brup3_2 | 25925 | 16191.44 ± 516.51 | 3.19 | 1.45 |  |
| Urkulu | Brup4_1 | 5165 | 17808.01 ± 498.62 | 2.8 | 1.41 | 1.407 ± 0.014 |
|  | Brup4_2 | 3150 | 17785.63 ± 492.66 | 2.77 | 1.42 |  |
|  | Brup5_1 | 3293 | 16792.42 ± 465.15 | 2.77 | 1.39 |  |
|  | Brup5_2 | 1084 | 17274.39 ± 452.59 | 2.62 | 1.4 |  |
|  | Brup6_1) | 17300 | 17384.86 ± 525.02 | 3.02 | 1.38 |  |
|  | Brup6_2 | 4333 | 16886.92 ± 538.69 | 3.19 | 1.41 |  |
