## Supplementary material for "Exploring genomic diversity and reproductive strategies in three expansion phases of the superdominant *Brachypodium rupestre* in high mountain grasslands": Table S4

**Table S4**. Ploidy level assignment of the studied samples through site-based heterozygosity using nQuack. Model selection and ploidy assignments. (A) Results of model selection to identify the most suitable model for ploidy estimation using as controls diploid (Brachypodium stacei, B. distachyon) and allotetraploid (B. rupestre, Arpea and Urkullu) individuals of known ploidy. The tested models include different distributions (Beta, Beta-Binomial, Beta-Uniform, and Normal) and fixed variance types (fixed, fixed_2, fixed_3). The column "correct" indicates the number of individuals assigned to the known expected ploidy (diploid or tetraploid) for each model, the best model (normal-uniform fixed) is highlighted in bold. (B) Ploidy assignment results for all analyzed individuals using the selected Normal-Uniform Fixed model. Control diploid individuals (B. stacei and B. distachyon, Campos et al. 2024) were included for validation, while the remaining individuals correspond to the B. rupestre samples under study. Each individual was assigned a ploidy level based on 1000 bootstrap replicates, confirming their expected diploid or tetraploid nature.

(A)

| **Distribution** | **Type** | **Total** | **Correct** |
| --- | --- | --- | --- |
| beta | fixed | 8 | 0 |
| beta | fixed_2 | 8 | 4 |
| beta | fixed_3 | 8 | 4 |
| beta-binomial | fixed | 8 | 0 |
| beta-binomial | fixed_2 | 8 | 4 |
| beta-binomial | fixed_3 | 8 | 4 |
| beta-binomial-uniform | fixed | 8 | 0 |
| beta-binomial-uniform | fixed_2 | 8 | 4 |
| beta-binomial-uniform | fixed_3 | 8 | 4 |
| beta-uniform | fixed | 8 | 0 |
| beta-uniform | fixed_2 | 8 | 4 |
| beta-uniform | fixed_3 | 8 | 4 |
| normal | fixed | 8 | 4 |
| normal | fixed_2 | 8 | 4 |
| normal | fixed_3 | 8 | 4 |
| **normal-uniform** | **fixed** | **8** | **8** |
| normal-uniform | fixed_2 | 8 | 6 |
| normal-uniform | fixed_3 | 8 | 4 |

(B)

| **Species** | **Ind** | **Model** | **Type** | **Bootstrap** | **Inference** |
| --- | --- | --- | --- | --- | --- |
| *B. stacei* | Canaries1 | Normal Uniform | Fixed | 859 | Diploid |
| *B. stacei* | Jaen2 | Normal Uniform | Fixed | 1000 | Diploid |
| *B. distachyon* | Adi2 | Normal Uniform | Fixed | 963 | Diploid |
| *B. distachyon* | Tek2 | Normal Uniform | Fixed | 1000 | Diploid |
| *B. rupestre* | Arpea-I1_1 | Normal Uniform | Fixed | 1000 | Tetraploid |
| *B. rupestre* | Arpea-I1_2 | Normal Uniform | Fixed | 1000 | Tetraploid |
| *B. rupestre* | Arpea-I1_3 | Normal Uniform | Fixed | 1000 | Tetraploid |
| *B. rupestre* | Arpea-I1_4 | Normal Uniform | Fixed | 1000 | Tetraploid |
| *B. rupestre* | Arpea-I1_5 | Normal Uniform | Fixed | 1000 | Tetraploid |
| *B. rupestre* | Arpea-I1_6 | Normal Uniform | Fixed | 1000 | Tetraploid |
| *B. rupestre* | Arpea-I1_7 | Normal Uniform | Fixed | 1000 | Tetraploid |
| *B. rupestre* | Arpea-I1_8 | Normal Uniform | Fixed | 1000 | Tetraploid |
| *B. rupestre* | Arpea-I1_9 | Normal Uniform | Fixed | 1000 | Tetraploid |
| *B. rupestre* | Arpea-I1_10 | Normal Uniform | Fixed | 1000 | Tetraploid |
| *B. rupestre* | Arpea-I1_11 | Normal Uniform | Fixed | 1000 | Tetraploid |
| *B. rupestre* | Errozate-I2_1 | Normal Uniform | Fixed | 1000 | Tetraploid |
| *B. rupestre* | Errozate-I2_2 | Normal Uniform | Fixed | 1000 | Tetraploid |
| *B. rupestre* | Errozate-I2_3 | Normal Uniform | Fixed | 1000 | Tetraploid |
| *B. rupestre* | Errozate-I2_4 | Normal Uniform | Fixed | 1000 | Tetraploid |
| *B. rupestre* | Errozate-I2_5 | Normal Uniform | Fixed | 1000 | Tetraploid |
| *B. rupestre* | Errozate-I2_6 | Normal Uniform | Fixed | 1000 | Tetraploid |
| *B. rupestre* | Errozate-I2_7 | Normal Uniform | Fixed | 1000 | Tetraploid |
| *B. rupestre* | Errozate-I2_8 | Normal Uniform | Fixed | 1000 | Tetraploid |
| *B. rupestre* | Errozate-I2_9 | Normal Uniform | Fixed | 1000 | Tetraploid |
| *B. rupestre* | Errozate-I2_10 | Normal Uniform | Fixed | 1000 | Tetraploid |
| *B. rupestre* | Errozate-I2_11 | Normal Uniform | Fixed | 1000 | Tetraploid |
| *B. rupestre* | Errozate-I2_12 | Normal Uniform | Fixed | 1000 | Tetraploid |
| *B. rupestre* | Errozate-P1_13 | Normal Uniform | Fixed | 1000 | Tetraploid |
| *B. rupestre* | Errozate-P1_14 | Normal Uniform | Fixed | 1000 | Tetraploid |
| *B. rupestre* | Errozate-P1_15 | Normal Uniform | Fixed | 1000 | Tetraploid |
| *B. rupestre* | Errozate-P1_16 | Normal Uniform | Fixed | 1000 | Tetraploid |
| *B. rupestre* | Errozate-P1_17 | Normal Uniform | Fixed | 1000 | Tetraploid |
| *B. rupestre* | Errozate-P1_18 | Normal Uniform | Fixed | 1000 | Tetraploid |
| *B. rupestre* | Errozate-P1_19 | Normal Uniform | Fixed | 1000 | Tetraploid |
| *B. rupestre* | Errozate-P1_20 | Normal Uniform | Fixed | 1000 | Tetraploid |
| *B. rupestre* | Errozate-P1_21 | Normal Uniform | Fixed | 1000 | Tetraploid |
| *B. rupestre* | Errozate-P1_22 | Normal Uniform | Fixed | 1000 | Tetraploid |
| *B. rupestre* | Errozate_7 | Normal Uniform | Fixed | 1000 | Tetraploid |
| *B. rupestre* | Ezkanda-S1_2 | Normal Uniform | Fixed | 1000 | Tetraploid |
| *B. rupestre* | Ezkanda-S1_3 | Normal Uniform | Fixed | 1000 | Tetraploid |
| *B. rupestre* | Ezkanda-S1_4 | Normal Uniform | Fixed | 1000 | Tetraploid |
| *B. rupestre* | Ezkanda-S1_5 | Normal Uniform | Fixed | 1000 | Tetraploid |
| *B. rupestre* | Ezkanda-S1_6 | Normal Uniform | Fixed | 1000 | Tetraploid |
| *B. rupestre* | Ezkanda-S1_8 | Normal Uniform | Fixed | 1000 | Tetraploid |
| *B. rupestre* | Ezkanda-S1_9 | Normal Uniform | Fixed | 1000 | Tetraploid |
| *B. rupestre* | Mendizar-I3_1 | Normal Uniform | Fixed | 1000 | Tetraploid |
| *B. rupestre* | Mendizar-I3_2 | Normal Uniform | Fixed | 1000 | Tetraploid |
| *B. rupestre* | Mendizar-I3_3 | Normal Uniform | Fixed | 1000 | Tetraploid |
| *B. rupestre* | Mendizar-I3_4 | Normal Uniform | Fixed | 1000 | Tetraploid |
| *B. rupestre* | Mendizar-I3_5 | Normal Uniform | Fixed | 1000 | Tetraploid |
| *B. rupestre* | Mendizar-I3_6 | Normal Uniform | Fixed | 1000 | Tetraploid |
| *B. rupestre* | Mendizar-I3_7 | Normal Uniform | Fixed | 1000 | Tetraploid |
| *B. rupestre* | Mendizar-I3_8 | Normal Uniform | Fixed | 1000 | Tetraploid |
| *B. rupestre* | Mendizar-I3_9 | Normal Uniform | Fixed | 1000 | Tetraploid |
| *B. rupestre* | Mendizar-I3_10 | Normal Uniform | Fixed | 1000 | Tetraploid |
| *B. rupestre* | Mendizar-I3_11 | Normal Uniform | Fixed | 1000 | Tetraploid |
| *B. rupestre* | Mendizar-I3_12 | Normal Uniform | Fixed | 1000 | Tetraploid |
| *B. rupestre* | Mendizar-P2_1 | Normal Uniform | Fixed | 1000 | Tetraploid |
| *B. rupestre* | Mendizar-P2_2 | Normal Uniform | Fixed | 1000 | Tetraploid |
| *B. rupestre* | Mendizar-P2_3 | Normal Uniform | Fixed | 1000 | Tetraploid |
| *B. rupestre* | Mendizar-P2_4 | Normal Uniform | Fixed | 1000 | Tetraploid |
| *B. rupestre* | Mendizar-P2_5 | Normal Uniform | Fixed | 1000 | Tetraploid |
| *B. rupestre* | Mendizar-P2_6 | Normal Uniform | Fixed | 1000 | Tetraploid |
| *B. rupestre* | Mendizar-P2_7 | Normal Uniform | Fixed | 1000 | Tetraploid |
| *B. rupestre* | Mendizar-P2_8 | Normal Uniform | Fixed | 1000 | Tetraploid |
| *B. rupestre* | Mendizar-P2_9 | Normal Uniform | Fixed | 1000 | Tetraploid |
| *B. rupestre* | Orion-S2_1 | Normal Uniform | Fixed | 1000 | Tetraploid |
| *B. rupestre* | Orion-S2_2 | Normal Uniform | Fixed | 1000 | Tetraploid |
| *B. rupestre* | Orion-S2_3 | Normal Uniform | Fixed | 1000 | Tetraploid |
| *B. rupestre* | Orion-S2_4 | Normal Uniform | Fixed | 1000 | Tetraploid |
| *B. rupestre* | Orion-S2_5 | Normal Uniform | Fixed | 1000 | Tetraploid |
| *B. rupestre* | Orion-S2_6 | Normal Uniform | Fixed | 1000 | Tetraploid |
| *B. rupestre* | Orion-S2_7 | Normal Uniform | Fixed | 1000 | Tetraploid |
| *B. rupestre* | Orion-S2_8 | Normal Uniform | Fixed | 1000 | Tetraploid |
| *B. rupestre* | Orion-S2_9 | Normal Uniform | Fixed | 1000 | Tetraploid |
| *B. rupestre* | Orion-S2_10 | Normal Uniform | Fixed | 1000 | Tetraploid |
| *B. rupestre* | Urkulu-P3_1 | Normal Uniform | Fixed | 1000 | Tetraploid |
| *B. rupestre* | Urkulu-P3_2 | Normal Uniform | Fixed | 1000 | Tetraploid |
| *B. rupestre* | Urkulu-P3_3 | Normal Uniform | Fixed | 1000 | Tetraploid |
| *B. rupestre* | Urkulu-P3_4 | Normal Uniform | Fixed | 1000 | Tetraploid |
| *B. rupestre* | Urkulu-P3_6 | Normal Uniform | Fixed | 1000 | Tetraploid |
| *B. rupestre* | Urkulu-P3_7 | Normal Uniform | Fixed | 1000 | Tetraploid |
| *B. rupestre* | Urkulu-P3_8 | Normal Uniform | Fixed | 1000 | Tetraploid |
| *B. rupestre* | Urkulu-P3_9 | Normal Uniform | Fixed | 1000 | Tetraploid |
| *B. rupestre* | Urkulu-S3_1 | Normal Uniform | Fixed | 1000 | Tetraploid |
| *B. rupestre* | Urkulu-S3_2 | Normal Uniform | Fixed | 1000 | Tetraploid |
| *B. rupestre* | Urkulu-S3_3 | Normal Uniform | Fixed | 1000 | Tetraploid |
| *B. rupestre* | Urkulu-S3_4 | Normal Uniform | Fixed | 1000 | Tetraploid |
| *B. rupestre* | Urkulu-S3_5 | Normal Uniform | Fixed | 1000 | Tetraploid |
| *B. rupestre* | Urkulu-S3_6 | Normal Uniform | Fixed | 1000 | Tetraploid |
| *B. rupestre* | Urkulu-S3_7 | Normal Uniform | Fixed | 1000 | Tetraploid |
| *B. rupestre* | Urkulu-S3_8 | Normal Uniform | Fixed | 1000 | Tetraploid |
| *B. rupestre* | Urkulu-S3_9 | Normal Uniform | Fixed | 1000 | Tetraploid |
| *B. rupestre* | Urkulu-S3_10 | Normal Uniform | Fixed | 1000 | Tetraploid |
