## Supplementary material for "Exploring genomic diversity and reproductive strategies in three expansion phases of the superdominant *Brachypodium rupestre* in high mountain grasslands": Table S5

**Table S5**. Pairwise Fst values for genomic differentiation between populations of Brachypodium rupestre under study.

|  | **Errozate_I2** | **Errozate_P1** | **Mendizar_I3** | **Mendizar_P2** | **Orion_S2** | **Urkulu_S3** | **Urkulu_P3** | **Arpea_I1** | **Ezkanda_S1** |
| --- | --- | --- | --- | --- | --- | --- | --- | --- | --- |
| **Errozate_I2** | - | - | - | - | - | - | - | - | - |
| **Errozate_P1** | 0.1393 | - | - | - | - | - | - | - | - |
| **Mendizar_I3** | 0.1229 | 0.1691 | - | - | - | - | - | - | - |
| **Mendizar_P2** | 0.1060 | 0.1507 | 0.1251 | - | - | - | - | - | - |
| **Orion_S2** | 0.1381 | 0.1845 | 0.1634 | 0.1450 | - | - | - | - | - |
| **Urkulu_S3** | 0.1215 | 0.1681 | 0.143 | 0.1208 | 0.1540 | - | - | - | - |
| **Urkulu_P3** | 0.1299 | 0.1795 | 0.1567 | 0.1341 | 0.1645 | 0.1234 | - | - | - |
| **Arpea_I1** | 0.1172 | 0.1586 | 0.1419 | 0.1191 | 0.1501 | 0.1261 | 0.1326 | - | - |
| **Ezkanda_S1** | 0.1031 | 0.1549 | 0.1374 | 0.1087 | 0.1407 | 0.11745 | 0.1155 | 0.0987 | - |
